# Targeting Semaphorin 7a signaling in preclinical models of estrogen receptor-positive breast cancer

**DOI:** 10.1101/2025.05.21.655360

**Authors:** Rachel N Steinmetz, Veronica Wessells, Heather Fairchild, Traci R Lyons

**Author notes:** Corresponding author: Name: Traci R. Lyons Address: U. Colorado Anschutz, 12801 E 17^th^ Ave, L18-8103 MS 8117, Aurora, CO 80045.

## Abstract

Estrogen receptor-positive (ER+) breast cancer (BC) comprises over 70% of breast cancers and is the leading cause of BC-related deaths in women worldwide. Despite available therapies targeting ER in BC, recurrence occurs in many patients due to therapeutic resistance. Semaphorin 7a (SEMA7A) is a biomarker associated with poor prognosis and endocrine therapy resistance for BC patients. Survival analyses of ER+ BC patients on endocrine therapy confirm early recurrence in patients with SEMA7A+ tumors. Thus, we aim to establish novel treatment strategies to improve outcomes for patients with ER+ SEMA7A+ BC. In this paper we investigate the mechanisms by which SEMA7A promotes resistance to endocrine therapy and its potential as a therapeutic target for ER+ BC. Our studies suggest that SEMA7A binds to integrins β1 and β4 and activates AKT mediated pro-survival signaling via its RGD domain. Using syngeneic ER+ models, tumors in were treated with PI3K inhibitors (20mg/kg alpelisib; 10mg/kg GCT-007), alone or in combination with tamoxifen (0.5 mg/100uL peanut oil), which resulted in decreased tumor growth. Tumors were also treated with a combination of an anti-SEMA7A antibody (SmAbH1) (100-250mg/kg) and fulvestrant (83mg/kg), compared to single agents. Our results demonstrate that direct inhibition of SEMA7A via SmAbH1 significantly reduces tumor growth of SEMA7A+ tumors, and the combination with fulvestrant may be even more effective. Our studies suggest that patients with ER+SEMA7A+ tumors should be candidates for PI3K-targeted therapies or anti-SEMA7A-based therapy.

**TRANSLATIONAL RELEVANCE:** Our studies propose inhibition of a novel target in estrogen receptor-positive (ER+) breast cancer (BC) with an anti-Semaphorin 7a (SEMA7A) monoclonal antibody (SmAbH1). We show efficacy of SmAbH1 as a single agent and in combination with endocrine therapy in preclinical models. We also show that targeting of SEMA7A signaling, including the use of well-known and novel PI3K inhibitors, could be an effective treatment strategy for patients with SEMA7A+ BC, particularly in combination with standard of care. Our mechanistic studies detail the cellular signaling pathway activated by SEMA7A and support the further investigation of these novel drug combinations for clinical use. Further, SEMA7A is a biomarker for poor prognosis and decreased patient survival, posing the need for a clinical diagnostic test for SEMA7A levels in BC patients, which we are currently developing.

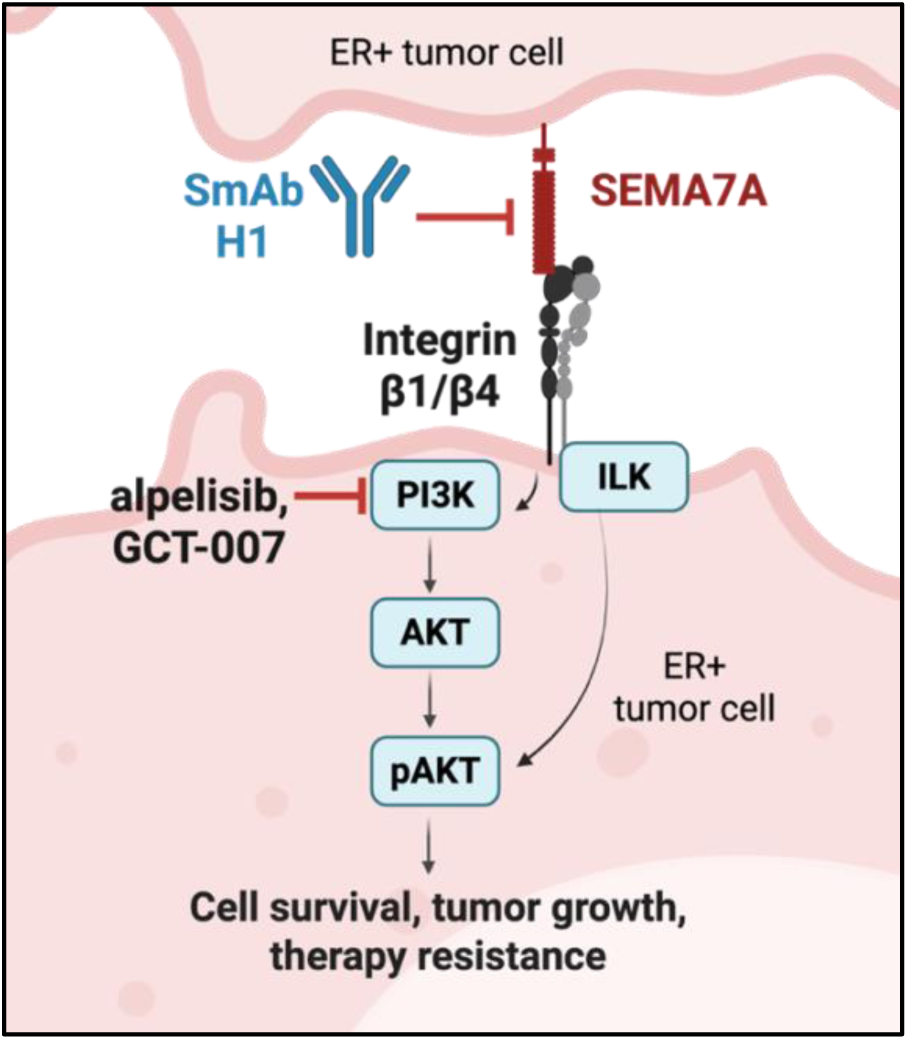

## INTRODUCTION

The most prevalent type of breast cancer (BC) is estrogen receptor-positive (ER+), accounting for 70% of all BC diagnoses (1,2). The standard treatment for women with ER+ BC typically involves endocrine therapy, such as the selective ER degrader, fulvestrant, the selective ER modulator, tamoxifen, and aromatase inhibitors such as exemestane and letrozole (2,3). While endocrine therapy is effective in many patients, 22-52% of patients with ER+ BC will experience disease recurrence, depending on lymph node involvement (2,3). Thus, there is an urgent need to improve treatments for patients with ER+ BC. We and others have identified that Semaphorin 7a (SEMA7A), a neuro-endocrine signaling protein (4), is dysregulated in various diseases and cancer types, resulting in poor prognosis (5–13). While numerous studies have investigated the role of SEMA7A in triple-negative breast cancer (TNBC), which lack expression of ER (ER-), progesterone receptor (PR-), and amplification of HER2 (HER2-) (14–19), we have shown that ER can regulate expression of SEMA7A. As such, and have previously analyzed the importance of SEMA7A expression in ER+ BC to show that SEMA7A is expressed in ER+ BC where expression confers resistance to ER targeting therapies including fulvestrant and tamoxifen (2). Our previous studies have also demonstrated mechanisms of therapy resistance activated in ER+ SEMA7A+ tumors that drive reduced sensitivity to the endocrine therapy fulvestrant, including downregulation of ER, activation of pro-survival signaling through Bcl-2, and regulation of markers of drug-resistant stem cells (CD44+/CD24-) (2). We have also shown that long-term estrogen deprivation, such as that induced by endocrine therapies, results in upregulation of SEMA7A and suggest pro-survival signaling as a potential mechanism of resistance(2). In support of this, other studies have shown that pro-survival signaling, such as that mediated by phosphoinositide linked kinase-3 (PI3K), AKT, and other protein kinases, leads to mechanisms of resistance to endocrine therapies (20,21); the contribution of PI3K/AKT to SEMA7A induced phenotypes is still unknown. Therefore, understanding how SEMA7A+ ER+ BCs upregulate pro-survival signaling, and whether these mechanisms could be targetable in SEMA7A+ tumors, will provide insight into treatment strategies for this subset of patients.

In previous results, we observed that *SEMA7A* mRNA expression is elevated in 67% of luminal breast cancer cell lines, compared to 50% and 36% in basal A and basal B, respectively (2) and that increased expression results in poor outcomes in all types of breast cancer(2,16). In its immune-related and other roles, SEMA7A is known to bind to two receptors: Plexin C1 and integrin β1 (22), and in models of TNBC we have shown that the pro-tumor roles for SEMA7A are dependent on integrin β1 and integrin α6 (14,16). However, in the context of ER+ BC, whether SEMA7A binds integrin receptors remains unknown. Known downstream targets of integrins are those involved in cell adhesion and cell survival, including the PI3K/AKT pathway. In support of a role for SEMA7A-and integrin-mediated activation of the PI3K pathway, studies in TNBC show activation of this pro-survival pathway (15). We have also shown that ER+SEMA7A+ tumors are sensitive to a non-selective PI3K inhibitor (LY294002) that is not utilized clinically. Clinical trials have also shown efficacy of alpelisib or alpelisib plus fulvestrant in PIK3CA-mutated ER+ breast cancer (23,24). Therefore, analysis of clinically relevant PI3K targeted therapies could provide insight into treatment options for patients with SEMA7A+ BC (2,25).

Constitutively active PI3K signaling can lead to worse patient prognosis and multidrug resistance, including fulvestrant resistance (20,26,27). We therefore sought to investigate the hypothesis that PI3K inhibitors could be effective for ER+SEMA7A+ BC. In our results presented herein, we show that breast cancer progression is increased in women with ER+SEMA7A+BC, resulting in poor prognosis and highlighting the need to identify additional treatment options. Mechanistically, we show for the first time that SEMA7A binds to integrin β1 and β4, in ER+ breast cancer cells and activates PI3K signaling as well as other phospho-signaling pathways related to cell survival in an integrin-dependent manner. Integrins can activate AKT directly through Integrin-linked kinase (ILK) phosphorylation of AKT at serine 473 (28) and we have previously published that inhibitors of PI3K, ILK, and/or integrins can reduce SEMA7A-mediated activation of pAKT (29); however, integrin and ILK inhibitors are not clinically available, so we explored the effect of PI3K inhibitors in ER+ SEMA7A+ tumors. We reveal a partial reversal of endocrine therapy resistance in ER+SEMA7A+ models of breast cancer by PI3K inhibition. Further, we tested a novel anti-SEMA7A monoclonal antibody for specifically reducing tumor growth in mice harboring SEMA7A+ tumors, both as a single agent and combined with fulvestrant. Collectively, our results support targeting of SEMA7A, both directly and indirectly through PI3K inhibition, as novel therapeutic strategies to improve patient outcomes in ER+SEMA7A+BC.

### MATERIALS & METHODS

#### Cell Culture

MCF7 cells were obtained from K. Horwitz (University of Colorado Anschutz Medical Campus). ZR75-1 cells were obtained from the University of Colorado Cancer Center Cell Technologies Shared Resource (CTSR). TC11 cells were obtained from L. Schuler (University of Wisconsin School of Veterinary Medicine). SSM2 cells were obtained from A. Borowsky (University of California Davis).

Cell lines were validated by fingerprinting analysis at the University of Colorado Anschutz/Barbara Davis Center Molecular Biology Core. Cells were routinely tested for mycoplasma contamination throughout the studies described herein with Lonza MycLonza MycoAlert PLUS (Walkersville, MD); last tested 3-24-2025. Cells were cultured at 37°C and 5% CO2. MCF7 were grown in MEM with 5% heat-inactivated fetal bovine serum (FBS; Peak Serum PS-FB3, Colorado USA) and 1% penicillin/streptomycin (P/S; ThermoFisher Scientific). TC11 and ZR75-1 were grown in RMPI 1640 with 10% heat-inactivated FBS and 1% P/S. SSM2 were grown in DMEM/F12 and supplemented with 10% FBS, 1% L-glutamine, 1% P/S. 50 μM 2-mercaptoethanol, 0.3 μM hydrocortisone, 5 μg/ml insulin and 10 ng/ml transferrin. Experiments were performed with cells below passage 30. For exogenous SEMA7A treatments, recombinant SEMA7A (rSEMA7A) was purified from SEMA7A-Fc expressing MDA-MB-231 cells in collaboration with the University of Colorado Cancer Center CTSR (14,30).

#### Transductions

MCF7-SEMA7A Overexpressing (OE) and MCF7 empty vector (EV) control cell populations were generated by lentiviral transductions with pLX304-Blast-V5 vectors; CMV-driven, V5-tagged full-length Semaphorin 7a cDNA (666 aa) or empty vector, respectively, and selected for using blasticidin at 5 μg/mL (Dana Farber Cancer Institute & CU Anschutz Functional Genomics Core). TC11-control and TC11-SEMA7A knockdown cell lines were generated via lentiviral transduction with pLKO-shNonTargeting-puro, and pLKO-shSEMA7A-038 (KD#1) and pLKO-shSEMA7A-040 (KD#2), respectively, and selected for using puromycin at 2 μg/mL. SEMA7A expression was validated via western blot analysis for all transduced cell lines.

#### Cell Viability Assays

Cells were plated in a 96-well (1,000 cells/well), 24-well (100,000 cells/well), 12-well (200,000 cells/well) or 6-well (500,000 cells/well) plate in triplicate and allowed to adhere at 37°C for 24 hours. After 24 hours, the cells were treated with vehicle or drug as follows: Fulvestrant and 4-hydroxytamoxifen were purchased from Millipore Sigma (Cat#129453-61-8 and Cat#68392-35-8, St. Louis, MO). For *in vitro* assays, fulvestrant and 4-hydroxytamoxifen treatments were at a final concentration of 8 nM and 5 μM in DMSO, respectively. Alpelisib (BYL719) was purchased from SelleckChem (Cat# S2814) and used at 0.5-1 μM in DMSO, as determined by our dose response and the literature (31). GCT-007 was obtained via collaboration with Global Cancer Technology (San Diego, CA) and used at 0.1-0.5 μM in DMSO as determined by our dose response. Anti-Semaphorin 7a monoclonal antibody; clone H1 (SmAbH1) was obtained from the University of Colorado Cancer Center CTSR. SmAbH1 or Mouse IgG1 (Bio X Cell; Cat# BP0083) was used at 1 μM in PBS as determined by our dose response and the literature (29,32,33). Percent confluence was determined via formalin fixation and 0.1% crystal violet staining and analyzed via ImageJ analysis. Data are normalized to control condition for each treatment. Experiments were performed in biological and technical triplicates and statistics were measured via repeated-measures two-way ANOVA with multiple comparisons.

#### Drug Titration Studies

5,000 SEMA7A overexpressing or empty vector control MCF7 cells, or 10,000 TC11 shCtrl or shSEMA7A cells, were seeded per well in a 96-well plate and allowed to adhere overnight. Vehicle or increasing concentrations of drug were added. Growth was assessed using 0.1% crystal violet staining after 24 hours of treatment. To calculate IC50 values, cell viability data for a range of concentrations (minimum 6-points data curve) for each drug was plotted as a dose-response curve in GraphPad Prism and analyzed using nonlinear regression three-parameters curve fit model of log(inhibitor) vs. response. Experiments were performed in triplicate and two-tailed, unpaired t-test was performed on each IC50 value in triplicate experiments.

#### Co-immunoprecipitation

Pierce Classic Magnetic IP/Co-IP Kit (ThermoFisher; Cat# 88804) and User Guide/CoIP Protocol (Pierce Biotechnologies) was used to perform all CoIP experiments. Cells were cultured normally to optimal confluence in T75 flasks. For RGDS experiments, RGDS peptide (Tocris Bioscience; Cat#3498, 50uM in milliQ H_2_O) or control (milliQ H_2_O) was added to culture media for 1 hour prior to making cell lysates. Lysates were made using IP lysis/wash buffer and incubated on ice for 15 min followed by centrifugation. Protein concentration of lysates was quantified using Qubit Protein Broad Range Assay Kit (ThermoFisher; Cat# A50668). Per sample, 1.0 mg of lysate was pre-incubated with 30 uL IP antibody: mouse IgG or SEMA7A (Santa Cruz; Sc-374432) and rocked overnight at 4°C. Per sample, 25μl of Pierce Protein A/G magnetic beads were washed and incubated with the antibody-lysate solution and rocked for 1 hour at room temperature. Using a magnet, supernatant was removed, and the beads were washed with lysis buffer three times. Beads were then eluted with Pierce Elution Buffer, and 10uL (pH) Neutralization Buffer was added per sample. Non-reducing Lane Marker Sample Buffer (5X, Pierce) was used as loading buffer for immunoblots following CoIP elution.

#### Immunoblotting

Protein extracts were prepared from cell lines by lysing cells in RIPA with EDTA (Thermo Fisher; Cat# J61529.AP) containing PhosSTOP phosphatase inhibitor (Roche; Cat# 4906837001), and complete protease inhibitor (Roche; Cat# 11697498001). Protein concentration of lysates was determined using Qubit Broad Range Protein Assay Kit (ThermoFisher; Cat# A50668). 30μg of total protein in Laemmli protein sample buffer was heated for 10 minutes. Lysates were run on SDS-Page gels and transferred onto methanol-activated PVDF membranes using the Trans-Blot Turbo Transfer System (Bio-Rad).

Membranes were blocked with 5% bovine serum albumin and probed with the following primary antibodies according to manufacturer recommendation: SEMA7A (Santa Cruz; Cat#Sc-374432), pAKT S473 (Cell Signaling; Cat# 4060), total AKT (Cell Signaling; Cat#2920), Integrin β4 (Cell Signaling; Cat#14803), Integrin β1 (Cell Signaling; Cat#34971), ERα (AbCam; Cat#16660), α-tubulin (Cell Signaling; Cat +#9099S) and GAPDH (Biolegend; Cat# 607902). Membranes were washed with 1X Tris-buffered saline with Tween 20, incubated with a goat anti-rabbit HRP secondary antibody (Abcam; Cat# 6721), and developed with enhanced luminescence detection system (Thermo Fisher; Cat# 32106) on the Licor Odyssey. Densitometric O.D. values of three biological replicates were obtained using ImageJ(34) and normalized to the loading control GAPDH or α-tubulin, and normalized to the first sample.

#### Animal Experiments

Power calculations based on pilot studies were utilized to determine the number of animals per group. Tumors were measured using digital calipers every 2–3 days with the observer blinded to treatment group, and tumor volume was calculated as (L × W × W × 0.5). Tumors were harvested at designated study endpoints (Maximum tumor volume = 1000mm^3^). Mammary glands with intact tumor, axial lymph nodes and lungs were harvested, formalin fixed, and paraffin embedded for immunohistochemical analysis. All animal studies were performed under the approval of the UC Anschutz Medical Campus Institutional Animal Care and Use Committee. Beeswax estrogen (E2) and sham pellets were generated and administered based on previous literature (35). For the study that combined fulvestrant and rSmAbH1-P, tumors were harvested and detached from the mammary gland; photographs were taken next to ruler and scaled to 1cm scale bar.

##### MCF7 Model

MCF7 is a well-known human ER+ cell line used to study breast cancer, thus it was an appropriate model to initiate our studies. 2,500,000 SEMA7A overexpressing or control MCF7 cells were injected into the #4 mammary fat pads of 6–8-week-old female NCG mice (Cat# CRL572, Charles River, Wilmington, MA). For drug treatment studies, fulvestrant dosing began when average tumor volume reached ∼100mm^3^. At day 37, animals were randomized to treatment into size-matched groups to receive either vehicle or PI3K inhibitor (alpelisib or GCT-007) daily.

##### TC11 Model

TC11 is a syngeneic ER+ murine model which we chose to use because it provides the context of an intact immune system, and there are limited other ER+ mouse models. TC11 also expresses endogenous SEMA7A, making it a relevant model for our research. 200,000 TC11 cells (wild type, shCtrl or shSEMA7A #2) were injected into the bilateral #4 mammary fat pads of 6–8-week-old female FVB/N mice (FVB/NCrl, Cat# CRL207, Charles River, Wilmington, MA). Mice were randomized to treatment into size-matched groups at average tumor volume ∼50-100mm^3^. For the estrogen/sham pilot study, mice were randomized to groups receiving E2 or sham pellets two days prior to tumor injections.

##### SSM2 Model

SSM2 is another syngeneic ER+ murine model which we chose to use because it provides the context of an intact immune system, and there are limited other ER+ mouse models. SSM2 also expresses endogenous SEMA7A, making it a relevant model for our research. 1,000,000 SSM2 tumor cells were injected into the #4 mammary fat pads of 6–8-week-old female 129 Sv-E mice (129 Sv-E, Cat# CRL287, Charles River, Wilmington, MA). Mice were randomized to treatment into size-matched groups at average tumor volume ∼50mm^3^.

##### Treatments

Fulvestrant was administered as follows: 2.5 mg in 90% sesame oil and 10% ethanol, injected subcutaneously every 5 days. This dose is equivalent to approximately 83 mg/kg per mouse. Using the FDA Dosing Guidance for Industry: Conversion of Animal Doses to Human Equivalent Doses, equivalent to 6.7mg/kg in humans (2). Tamoxifen was given using the following dosing regimen: 0.5 mg/100uL peanut oil per mouse, via intraperitoneal (IP) injection every 3^rd^ day (36). Alpelisib was dissolved in 1% Carboxymethyl cellulose solution + 0.5% Tween 80 and administered via oral gavage at 20 mg/kg daily. GCT-007 was dissolved in 55% 1M NaCl, 30% PEG-400, 7.5% Solutol-HS + 7.5% N-Methyl-2-pyrrolidone and administered via oral gavage at 10 mg/kg daily. SmAbH1/rSmAbH1 or Mouse IgG1 (Bio X Cell; Cat# BP0083) was given in sterile PBS at 100-250 μg per 100μL PBS every other day via IP injection (32,33).

#### Immunohistochemistry

Tissue collected from the *in vivo* study endpoints as described above, was formalin-fixed and paraffin-embedded. Tissue was stained with Hematoxylin and Eosin. Four-μm sections were pretreated with 10X Dako Target Retrieval System. Slides were prepared in a Dako Autostainer with primary antibody (Integrin β4: Cell Signaling Cat#14803, SEMA7A AbCam Cat # ab255602, 9EG7/CD29: BD Pharmingen rat-anti-mouse CD29 clone 9EG7 Cat# 550531), followed by EnVision+ HRP Rabbit (Agilent Technologies cat# K4003) antibody for 30 mins. For 9EG7 staining, rat-on-mouse HRP polymer detection kit (Biocare cat# RT517H) was used for secondary. The tissue was incubated with chromogen 3,3’diaminobenzidine (Vector Labs catalog #SK-4105) for 10 minutes, and then the tissue was counter-stained with hematoxylin for 1 minute (Vector Labs catalog #H-3401-500). Stained images were acquired using an Olympus Fluorescent Microscope IX83. Quantification was performed using Cell Sense Analysis.

#### ELISA

ELISA Buffer Kit (Thermo Cat #CNB0011) was used to perform an indirect ELISA where the plate was coated with SEMA7A protein and probed with serial dilution concentrations of rSmAbH1-P to determine binding specificity. A 96-well plate was pre-coated with 5ug/well of purified SEMA7A (50ug/mL) protein in Coating Buffer A and incubated at 4 degrees overnight. Plate was washed with 1X wash buffer and then probed with primary antibody (rSmAbH1-P) at serial dilution concentrations, from 20 to 0 ug/mL, and incubated at 4 degrees overnight. Plate was washed with 1X wash buffer three times and then probed with secondary antibody (Goat anti-mouse IgG-HRP; Abcam Cat#ab97040) for 30 minutes at room temperature. Plate was washed three times with 1X wash buffer and absorbance at 425nm was read on plate reader.

#### Data Mining Analysis

Kaplan Meir Plotter(37) was used to assess overall survival (OS), distant metastasis-free survival (DMFS), and post-progression survival (PPS) data for BC patients in the ER+ dataset, and patients in the ER+ dataset who received endocrine therapy (tamoxifen or fulvestrant). SEMA7A was queried and stratified into high and low-expression groups using the auto-select best cutoff level function. The generated data, plots, hazard ratio (HR), and LogRank P value were downloaded from KMplotter.com. TCGA Human Data Analysis: We analyzed The Cancer Genome Atlas (TCGA) Breast Cancer study for the mRNA expression of Semaphorin 7a (*SEMA7A),* integrin β1 *(ITGB1), and* integrin β4 *(ITGB4)* collected from patient samples processed by RNAseq – IlluminaHiSeq. Gene expression analysis was performed using the Xena Functional Genomics Explorer(38). The log2(norm_count+1) value for mRNA expression was used to graph correlations between SEMA7A and the other genes in the TCGA Breast Cancer dataset. Each data was analyzed using Pearson’s correlation (rho) and p-values are indicated on each plot.

#### BioSpa Assay

500 MCF7 cells per well were plated in 96 well plates and incubated overnight to allow for adherence. Cells were then treated with 1μM IgG1 or 1μM SmAbH1 and immediately placed in a BioSpa Live Cell Imager (Agilent) for 4 days, with images of each well being taken every 4 h, at ×10 magnification. The cell count of each well was measured via brightfield count in the BioSpa Software.

#### Reverse Phase Protein Array (RPPA)

For cell lysates (RPPA-1), MCF7 Empty Vector control or SEMA7A Overexpressing cells were cultured, each in quadruplicate, in a 10cm dish for 72 h. After washing cells with PBS, lysates were generated using RPPA Lysis Buffer (Baylor College of Medicine Antibody-Based Proteomics Core) and frozen at -20°C. For tumor RPPA (RPPA-2), tumors were harvested and flash-frozen in liquid nitrogen at necropsy and stored at -80°C. Lysates/tumors were shipped on dry ice to the Baylor College of Medicine (BCM) Antibody-Based Proteomics Core (Houston, TX). Subsequent RPPA was performed by the core facility as outlined here (https://www.bcm.edu/academic-centers/dan-l-duncan-comprehensive-cancer-center/research/cancer-center-shared-resources/antibody-based-proteomics/protocols-workflow) (39). Normalized fold change for each antibody was calculated by (each signal value/AVG value of controls) and is depicted as log2(normalized fold change. Significance was determined by performing an unpaired two-tailed t-test between the fold change values for each antibody. Each sample was run in technical triplicates and n=4 samples per group. Samples with a Coefficient of Variation >25% for technical triplicates were excluded from further analysis. Pathway analysis was done using Gene Ontology Pathway Analysis.

#### Protein Ribbon Structures

The protein structure of SEMA7A was analyzed in PyMOL with an academic research license. Ribbon structure (purple) was created and modified to highlight the RGD motif (orange) and the antibody-binding site (green) in PyMOL.

#### Synergy Scores

Bliss Synergy scores for two-drug combination treatments were calculated using Synergyfinder.org; scores ≤ -10 are considered antagonistic, between -10 and 10 are considered additive, and a score ≥ 10 is considered synergistic (40). Further details on the methods used can be found at https://synergyfinder.fimm.fi/synergy/synfin_docs/.

#### Novel drug treatments

##### SmAbH1 (hybridoma)

Mice were injected with SEMA7A protein and KLH peptide to enhance immune response. B cells from mouse spleens were fused with myeloma cells to create a hybridoma; positive hybridoma clones were selected and monoclonal antibodies were harvested. SmAbH1 Antibody binds to amino acids 381-392 of SEMA7A (green; SFigure 1G) (29).

##### rSmAbH1 (recombinant)

The ExpiCHO-S expression system (Gibco) was used for the production of recombinant antibodies according to the manufacturer’s protocol. After the 12 days of expression, the culture media were harvested by centrifugation, filtered through a 0.2-micron filter, and underwent affinity chromatography using MabSelect PrismA resin and the AKTA pure FPLC instrument for purification of recombinant monoclonal antibody.

##### GCT-007

GCT-007 is a pan PI3K inhibitor and was provided to us for these studies by Global Cancer Technology. (16776 Bernardo Center Drive #203, San Diego, CA 92128).

#### Statistical Analysis

All quantitative and statistical analysis was performed using GraphPad Prism (Version 10.3.1), and the methods of tests are labeled in the figure legends. The statistical tests used in this paper include unpaired two-tailed t-test, Pearson’s correlation, and ordinary one-way ANOVA with Tukey’s multiple comparisons test. All p-values were measured under two-tailed conditions. Significance was determined by a p-value less than 0.05. *p<0.05, **p<0.01, ***p<0.001 ****p<0.0001.

#### Data Availability

The data generated in this study are available upon request from the corresponding author.

## RESULTS

### SEMA7A is expressed in ER+ breast cancer and associated with decreased 5-year patient survival

To test our hypothesis that SEMA7A predicts decreased response to therapy, and therefore poor prognosis, we used Kaplan-Meier Plotter(37) to examine 5-year survival for ER+ BC patients with high versus low tumor *SEMA7A* mRNA expression (Figure 1A-C). The probability of overall survival (OS) was significantly decreased for patients with high *SEMA7A* expression (Figure 1A), as well as distant metastasis-free survival (DMFS) (Figure 1B) and post-progression survival (PPS) (Figure 1C). Further, relapse-free survival (RFS) (Figure 1D) and DMFS (Figure 1E) were decreased in ER+ BC patients with high *SEMA7A* expression who received endocrine therapy.

**Figure 1.**
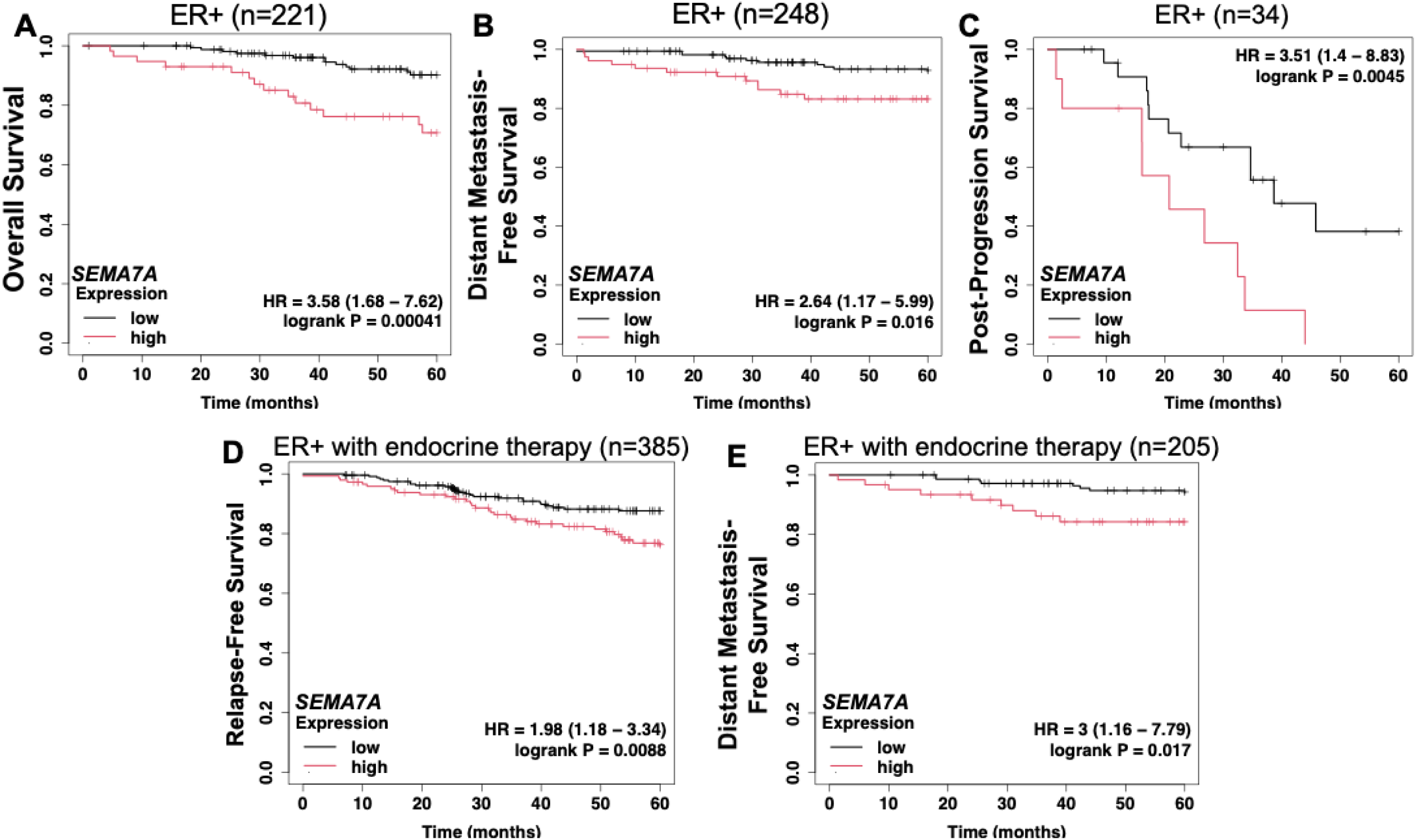
SEMA7A is expressed in ER+ breast cancer and is associated with decreased 5-year patient survival. Survival analysis was done via Kaplan-Meier Plotter, which was assessed for *SEMA7A* mRNA expression levels in breast cancer patients in the ER+ subtype dataset (n=3499). Data are shown as KM plots; curves are stratified by *SEMA7A* high (red) and low (black) expression. Hazard ratios (HR) and log-rank p-values are included. A) Overall survival; ER+ subtype (OS; n=221). B) Distant metastasis-free survival; ER+ subtype (DMFS; n=248). C) Post-progression survival; ER+ subtype (PPS; n=34). Further analysis was done within the ER+ subtype to include only patients who had received endocrine therapy (n=1831). D) Relapse-free survival; ER+ and received endocrine therapy (RFS; n=385). E) Distant metastasis-free survival: ER+ and received endocrine therapy (DMFS; n=205).

### SEMA7A activates AKT signaling and binds to Integrins β1 and β4 via RGD in ER+ tumor cells

To investigate differential protein and phospho-protein expression in ER+ MCF7 breast cancer cells transduced with an empty vector (EV) or SEMA7A overexpression (OE) vector, we utilized a reverse phase protein array (RPPA-1) (41). We confirmed SEMA7A overexpression via western blot prior to submission of lysates for RPPA (SFigure 1A). Using the data obtained from RPPA-1 we performed Gene Ontology (GO) Analysis to identify upregulated pathways in the SEMA7A OE lysates compared to EV; we observed that signaling related to cell survival, including AKT, mTOR, and integrin signaling, was upregulated in SEMA7A OE cells (Figure 2A). Specifically, the RPPA data suggest that overexpression of SEMA7A leads to increased integrins β1 and β4 as well as phospho-AKT (pAKT) at serine 473 (S473) (Figure 2B). The RPPA also show that integrin-linked kinase, which can activate AKT through S473 phosphorylation (42), the anti-apoptotic protein Bcl-2, and downstream targets of AKT, such as p-P70S6K (S371), were all increased in SEMA7A OE lysates (Figure 2B).

**Figure 2.**
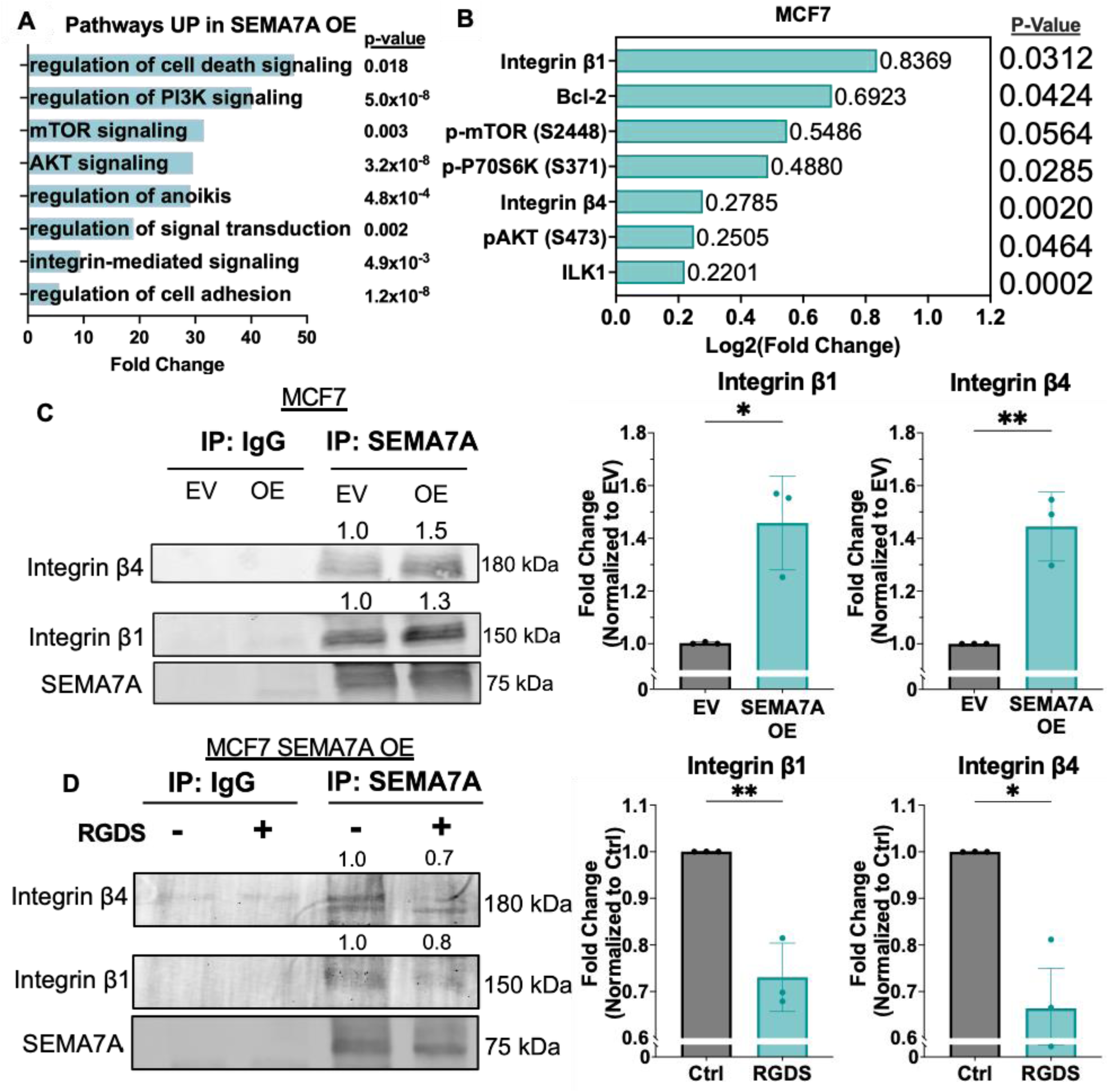
SEMA7A activates AKT signaling and binds to Integrins β1 and β4 via RGD in ER+ tumor cells. A) Gene Ontology (GO) Pathway Analysis of RPPA results, showing top pathways upregulated in MCF7 SEMA7A OE cell lysates, as fold change compared to EV, p<0.05. B) Log2(fold change) of integrin and AKT-related proteins in MCF7 SEMA7A OE versus EV lysates from RPPA-1 data. Multiple two-tailed unpaired t-tests. C) Immunoblot for Integrin β4, Integrin β1 and SEMA7A in elutions following CoIP of MCF7 EV and SEMA7A OE lysates for IgG or SEMA7A. ii. Quantification of IP shown as fold change compared to EV. Two-tailed unpaired t-test. D) Immunoblot for Integrin β4, Integrin β1 and SEMA7A in elutions following CoIP of MCF7 lysates for IgG or SEMA7A, after treatment with PBS (-) or 50 μM RGDS (+). Quantification of IP shown as fold change compared to control. Two-tailed unpaired t-test. Error bars are mean +/-SD. *p<0.05, **p<0.01, ***p<0.001, ****p<0.0001.

Phospho-mTOR(S2448), which is also a downstream target of AKT, was also increased, but not with statistical significance (p=0.0564)(Figure 2B). To validate that SEMA7A activates AKT signaling through pAKT (S473) in ER+ breast cancer cells, we validated pAKT (S473) and total AKT using immunoblot analysis on MCF7 EV and SEMA7A OE cells, or WT MCF7 treated with purified SEMA7A (rSEMA7A); the pAKT to AKT ratio was significantly higher in SEMA7A OE cells compared to EV and rSEMA7A-treated cells compared to controls (SFigure 1B-C). Expression of *ITGB1* and *ITGB4* mRNA also weakly correlated with *SEMA7A* expression in the TCGA Breast Cancer Dataset (n=1247) (SFigure 1D-E). To investigate integrin β1 and β4 as binding partners for SEMA7A in ER+ breast cancer cells, we performed co-immunoprecipitation (CoIP) using a SEMA7A specific antibody to bind the protein and its binding partner(s). After CoIP, western blotting revealed binding of both integrin β1 and β4 to SEMA7A, which was higher in SEMA7A OE cells compared to controls (Figure 2C, SFigure 1G: CoIP input and unbound controls). To investigate whether SEMA7A binds to integrins via its RGD (Arg-Gly-Asp) motif (SFigure 1F), which is known to facilitate integrin binding (43), we conducted CoIP using MCF7 SEMA7A OE lysates treated with PBS or RGDS (Arg-Gly-Asp-Ser) peptide, which inhibits RGD-integrin binding. Our results indicate that binding of SEMA7A to integrin β1 and β4 was significantly reduced by the RGDS peptide (Figure 2D, SFigure 1H: CoIP input and unbound contorls). These results were validated in an additional human ER+ breast cancer cell line, ZR75-1 (SFigure 1I).

### SEMA7A-mediated activation of AKT is dependent on RGD binding in ER+ tumor cells

Based on our results showing increased pAKT (S473) in SEMA7A OE MCF7, we validated our results in a syngeneic ER+ cell line. We obtained TC11 and SSM2 cells, which are ER+ mouse mammary tumor cell lines (44,45)(generous gifts from Linda Schuler and Alexander Borowsky), and validated ERα expression compared to MCF7 (SFigure 2A). We also compared SEMA7A expression in the two cell lines to reveal that TC11 cells express slightly higher endogenous SEMA7A (SFigure 2B). To determine whether SEMA7A was necessary to drive pAKT in the TC11 cell line, we compared cell lines stably transduced with shCtrl and shSEMA7A lentiviral vectors (shSEMA7A KD#1 and KD#2).

We confirmed SEMA7A expression was decreased in both KD vectors, with a larger decrease in KD#2 (SFigure 2C). As expected, knockdown of SEMA7A decreased pAKT levels, and the ratio of pAKT to AKT was decreased in a SEMA7A-dependent manner (Figure 3A). Additionally, we show that recombinant SEMA7A (rSEMA7A) or conditioned media from SEMA7A OE cells rescued the decrease in pAKT observed in the KD cells (Figure 3B). To further define whether AKT activation by SEMA7A occurs via SEMA7A-integrin binding, we treated TC11 cells with rSEMA7A, RGDS peptide or both, and measured pAKT/AKT expression (Figure 3C). Although the RGDS peptide alone inhibited pAKT levels, our results also show that rSEMA7A increased pAKT, and this effect was inhibited by the RGDS peptide (Figure 3C). We observed similar results with the RGDS in two additional ER+ cell lines, MCF7 and ZR751 (SFigure 2D-E).

**Figure 3.**
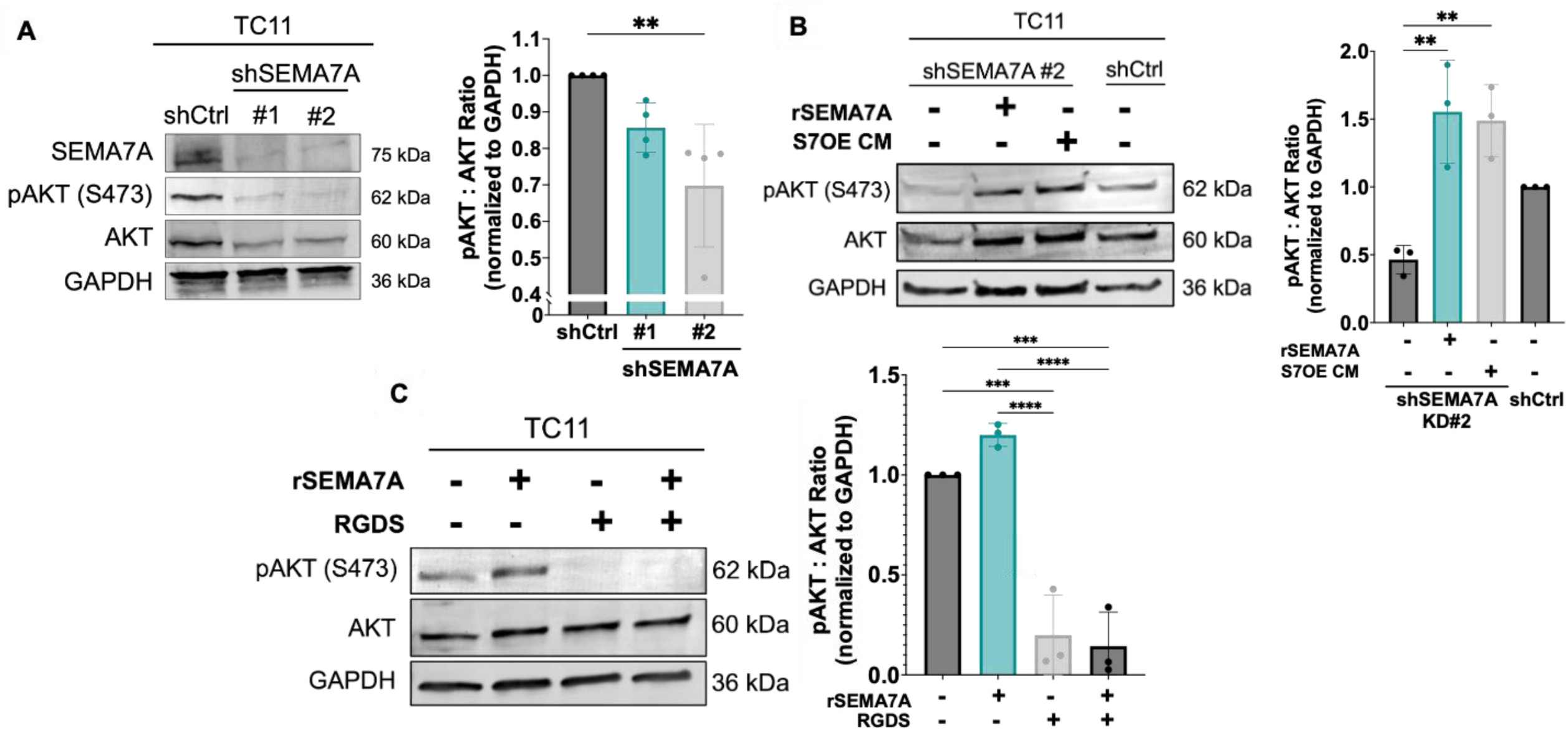
SEMA7A-mediated activation of AKT is dependent on RGD binding in ER+ tumor cells. A) Representative immunoblot for pAKT (S473), AKT and GAPDH in TC11 shCtrl and shSEMA7A (KD#1 and KD#2) cell lines. Bar graph: ratio of pAKT to total AKT. Ordinary one-way ANOVA with Tukey’s multiple comparisons test. B) Representative immunoblot for pAKT (S473), AKT and GAPDH in TC11 shCtrl and shSEMA7A KD#2 cell lines after treatment for 24 hours with 20 μg/mL purified rSEMA7A, or conditioned media from SEMA7A OE cells (S7OE CM). Bar graph: ratio of pAKT to total AKT. Ordinary one-way ANOVA with Tukey’s multiple comparisons test. C) Representative immunoblot for pAKT (S473), AKT and GAPDH in TC11 WT cell lysates after treatment for 1 hour with 20 μg/mL rSEMA7A, 50 μM RGDS peptide, or both. Bar graph: ratio of pAKT to total AKT. Ordinary one-way ANOVA with Tukey’s multiple comparisons test. Error bars are mean +/- SD. *p<0.05, **p<0.01, ***p<0.001, ****p<0.0001.

### SEMA7A+ tumor growth is inhibited by combination treatment with endocrine therapy and PI3K inhibitors

Since there are no clinically available integrin or ILK inhibitors, we investigated other therapeutic strategies to inhibit SEMA7A-mediated signaling. Increased pAKT levels in SEMA7A-expressing cells suggest a potential increase in PI3K signaling activity (46). Thus, we hypothesized that SEMA7A+ ER+ BC cells would exhibit increased sensitivity to treatment with PI3K inhibitors (PI3Ki).

To test this hypothesis, we determined the 50% inhibitory concentration (IC50) for alpelisib (p110α inhibitor) and GCT-007 (pan-PI3K inhibitor in development for treatment in glioblastoma and breast cancer(47)) in MCF7 EV and SEMA7A OE cells (SFigure 3A-B). The IC50 of alpelisib and GCT-007 in SEMA7A OE cells was lower, suggesting increased susceptibility to PI3Ki in SEMA7A-expressing tumor cells. We also examined sensitivity to PI3Ki in TC11 shSEMA7A #2 KD cells compared to controls (SFigure 3C-D). In support of our hypothesis, cell viability assays revealed that inhibition by both alpelisib and GCT-007 is greater in SEMA7A OE cells, compared to controls (SFigure 3E). We also confirmed that pAKT expression is reduced by both PI3Ki treatments in MCF7 and TC11 (SFigure 3F-G) cells.

Since PI3K inhibition is utilized clinically in conjunction with endocrine therapy for ER+ breast cancers, we aimed to evaluate whether a combination of endocrine therapy and PI3K inhibition (PI3Ki) would be more effective than either treatment alone in SEMA7A-expressing tumors. Based on previous literature (2) and our IC50 curves (SFigure 3A-D; SFigure 4A-B), we treated ER+ TC11 cells with fulvestrant, alpelisib, or the combination and measured cell viability to reveal that fulvestrant alone had minimal effect, while single-agent alpelisib and GCT-007 resulted in a moderate reduction in cell viability, and the combination treatments were the most effective (Figure 4A-B). Similar results were observed with the combination of fulvestrant plus alpelisib or GCT-007 in another ER+ cell line, ZR75-1 (SFigure 4C-D). Notably, TC11 tumor cells are reported to exhibit resistance to tamoxifen (44).

Therefore, we also tested the combination of tamoxifen and each PI3Ki in TC11 cells; our findings reveal that tamoxifen alone has minimal effect, whereas the combination significantly reduced cell viability compared to control, to a greater extent than either single agent (Figure 4C-D). Following *in vitro* synergy assays, Bliss synergy scores were calculated for the treatment combinations of fulvestrant (Figure 4E-F) and tamoxifen (Figure 4G-H) with PI3K inhibitors. Synergy scores between 0 and 10 indicate an additive effect, while scores above 10 indicate synergy between two treatments, and we observed synergy with fulvestrant/alpelisib (Score = 11.3), tamoxifen/alpelisib (Score = 14.61) and tamoxifen/GCT-007 (Score = 16.33), and an additive effect with fulvestrant/GCT-007 (Score = 3.52) (40). To evaluate the combination of endocrine therapy and PI3K inhibition *in vivo*, MCF7 SEMA7A OE tumor-bearing NCG mice were treated with fulvestrant alone or in combination with alpelisib or GCT-007, resulting in a greater reduction of tumor growth with the combination treatments (SFigure 4E-F). We monitored mouse weight and moribound criteria and observed no evidence of toxicities with either PI3K inhibitor based on these criteria (SFigure 4G-H). In data from the manufacturer, GCT-007 exhibits efficacy in immunocompetent mice using a model of glioblastoma (unpublished observations). Therefore, we tested combination treatments in an ER+ (SFigure 4I), estrogen-responsive (44)(SFigure 4J) immune-competent model. TC11 tumor-bearing FVB/N mice were treated with tamoxifen, GCT-007, or the combination; the combination was most effective at inhibiting tumor growth compared to either single agent or controls (Figure 4I). Consistent with the previous study, mouse no evidence of toxicity was observed (SFigure 4K).

**Figure 4.**
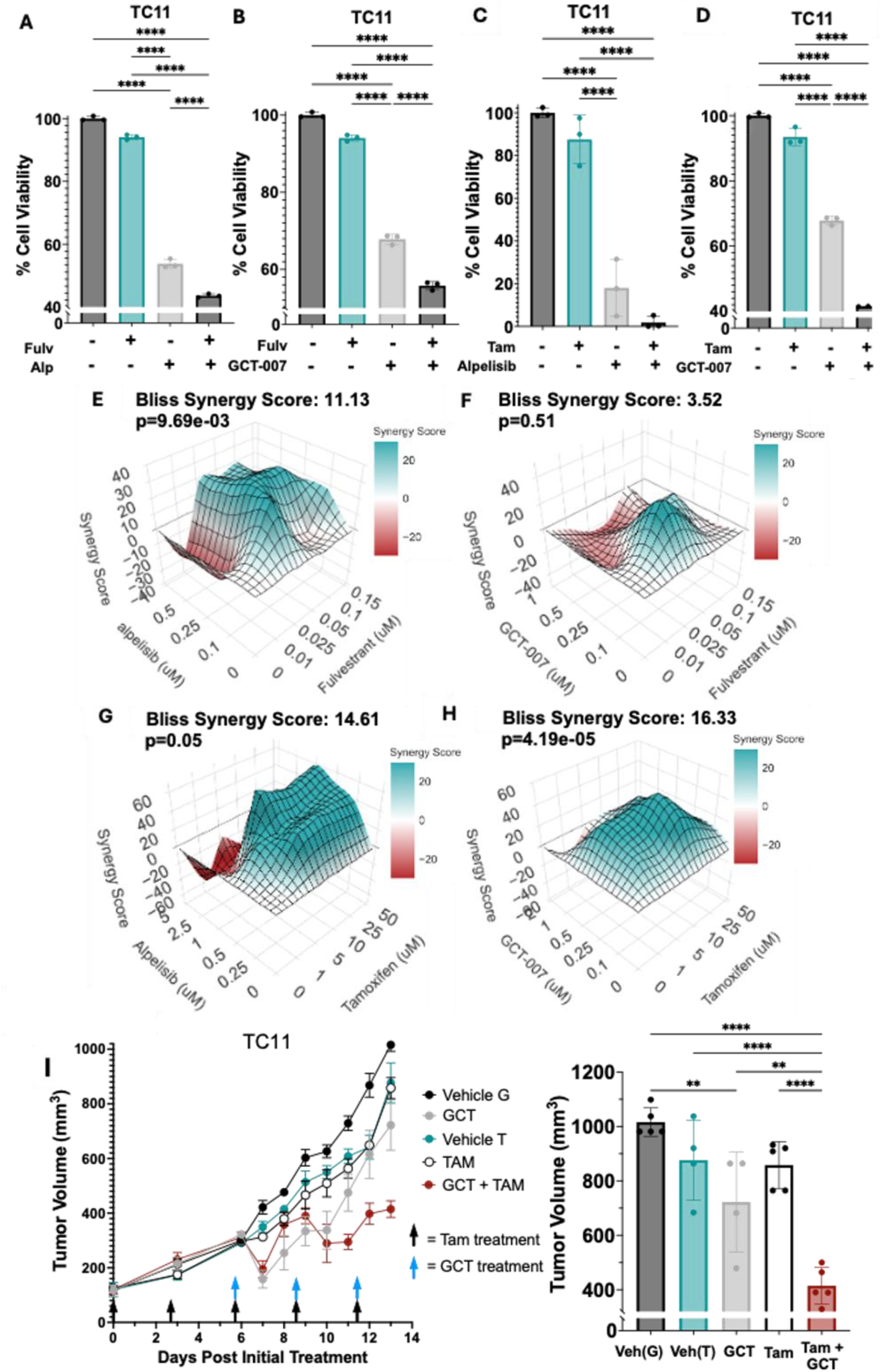
SEMA7A+ tumor growth is inhibited by combination treatment with endocrine therapy and PI3K inhibitors. A) Percent cell viability quantified via crystal violet staining after 48-hour treatment of TC11 cells with DMSO, 8nM fulvestrant, 1mM alpelisib, or the combination, normalized to control. Ordinary one-way ANOVA with Tukey’s multiple comparisons test. B) Percent cell viability quantified via crystal violet staining after 48-hour treatment of TC11 cells with DMSO, 8nM fulvestrant, 0.3mM GCT-007, or the combination, normalized to control. Ordinary one-way ANOVA with Tukey’s multiple comparisons test. C) Percent cell viability quantified via crystal violet staining after 48-hour treatment of TC11 cells with DMSO, 5 mM 4-hydroxytamoxifen, 1mM alpelisib, or the combination, normalized to control. Ordinary one-way ANOVA with Tukey’s multiple comparisons test. D) Percent cell viability quantified via crystal violet staining after 48-hour treatment of TC11 cells with DMSO, 5 mM 4-hydroxytamoxifen, 0.3 mM GCT-007, or the combination, normalized to control. Ordinary one-way ANOVA with Tukey’s multiple comparisons test. E) Bliss synergy score for fulvestrant and alpelisib in TC11 cells. F) Bliss synergy score for fulvestrant and GCT-007 in TC11 cells. G) Bliss synergy score for (4-hydroxy)tamoxifen and alpelisib in TC11 cells. H) Bliss synergy score for (4-hydroxy)tamoxifen and GCT-007 in TC11 cells. I) Average TC11 tumor volume over time in FVB/N mice, in groups treated with Vehicle, GCT-007, tamoxifen, or the combination. GCT = GCT-007, TAM = tamoxifen. Bar graph: tumor volume at the study endpoint (Day 13). Ordinary one-way ANOVA with Tukey’s multiple comparisons test. Error bars are mean +/- SD (A-D), or mean +/- SEM (I). *p<0.05, **p<0.01, ***p<0.001, ****p<0.0001.

### Inhibition of SEMA7A using a monoclonal antibody (SmAbH1) reduces tumor growth and AKT signaling in ER+ BC

We next tested our hypothesis that resistance to endocrine therapy, mediated by SEMA7A (2), could be reversed by direct targeting of SEMA7A using our novel anti-SEMA7A monoclonal antibody (SmAbH1) that we have previously shown to have efficacy in ER-negative models of breast cancer(32,33). Our *in vitro* BioSpa live imaging assay results show that cell viability is reduced by SmAbH1 over time in MCF7 SEMA7A OE cells, but not in EV controls (Figure 5A). The IC50 for SmAbH1 is also lower in SEMA7A OE compared to controls (SFigure 5A), suggesting SmAbH1 is most effective in SEMA7A-expressing cells. Previous studies from our lab have confirmed binding specificity of SmAbH1 to SEMA7A protein via ELISA (29). In contrast, TC11 SEMA7A shSEMA7A KD#2 showed less response to SmAbH1 compared to TC11 shCtrl cells *in vitro* (Figure 5B). Our previous studies suggest that SmAbH1 is not efficacious in immune-compromised hosts (32). Thus, we evaluated SmAbH1 in the TC11 model *in vivo* (SFigure 5B-D). We observed inhibition of tumor growth in SmAbH1-treated tumors only (SFigure 5B-C). We observed no evidence of toxicity based on mouse weight and moribound criteria (SFigure 5D). We validated these findings in the SSM2/129Sv-E model, an additional ER+ immune-competent model (48) with endogenous SEMA7A expression (SFigure 2B, SFigure 5E). As further evidence of specificity of SmAbH1, we observed that SmAbH1 inhibited tumor growth compared to the IgG control (Figure 5C) and this effect was specific to SEMA7A-expressing shCtrl tumors only (Figure 5C). Similarly, tumor volume calculated at study endpoint and tumor weight were significantly reduced only in the TC11 shCtrl tumors treated with SmAbH1 (Figure 5D-E). To confirm a SEMA7A/integrin-mediated mechanism *in vivo*, we performed IHC staining on control and SmAbH1-treated TC11 tumors (Figure 5F). To look at pAKT levels, we performed RPPA analysis (RPPA-2) of lysates made from IgG-treated and SmAbH1-treated TC11 and SSM2 tumors to reveal decreased expression of pAKT (S473) in SmAbH1-treated tumors (Figure 5G, SFigure 5F). Further, quantification of tumor IHC staining showed that activated integrin β1 (via 9EG7 staining), integrin β4, and SEMA7A protein expression were reduced in the SmAbH1-treated group (Figure 5H-J, SFigure 5G: secondary-only IHC controls, SFigure 5H-J: n=5 per group).

**Figure 5.**
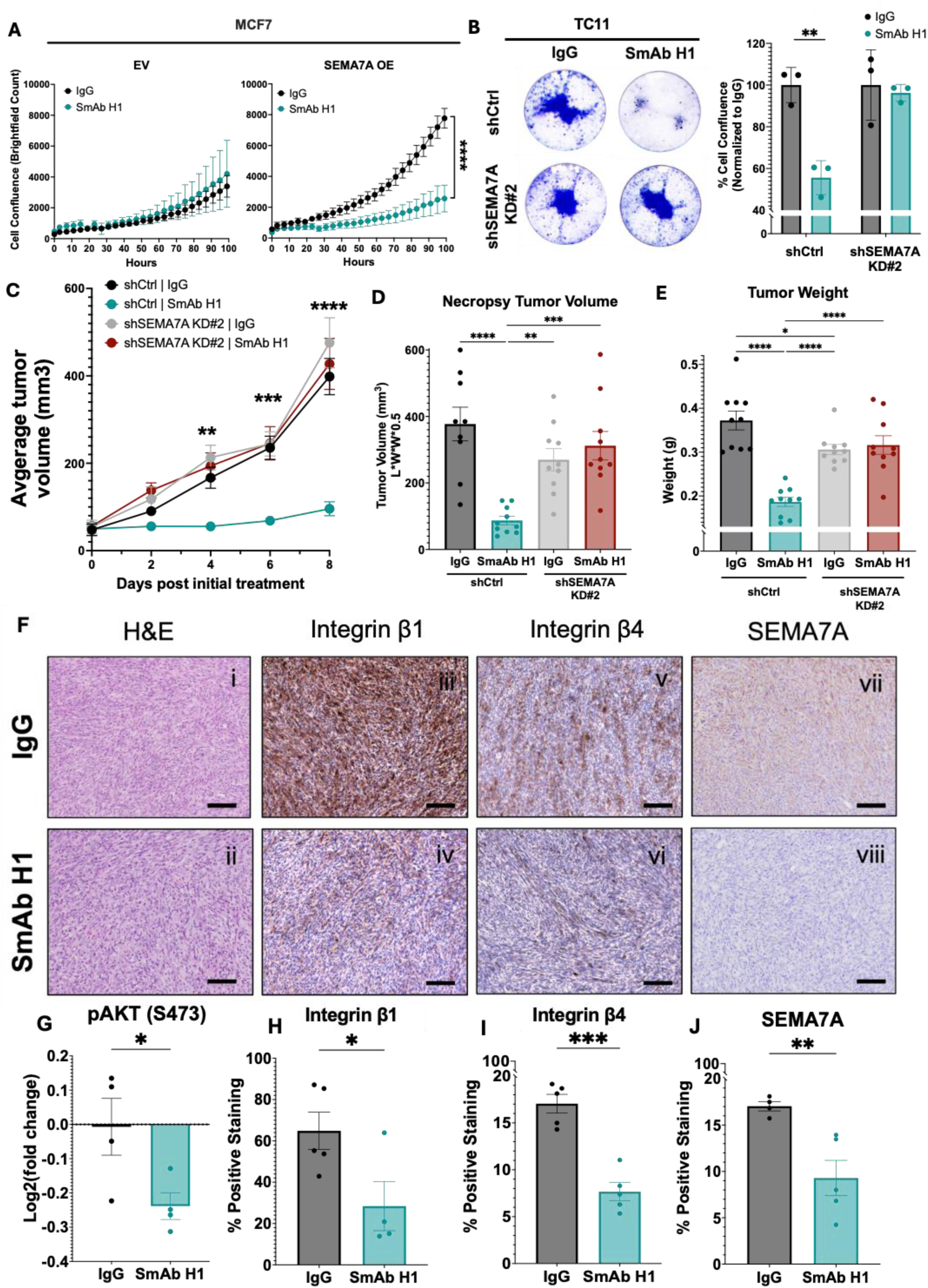
Direct inhibition of SEMA7A using a monoclonal antibody (SmAb H1) reduces tumor growth and AKT signaling in ER+ BC. A) Brightfield cell count measured by BioSpa Live Imaging over 96 hours of MCF7 EV and SEMA7A OE cells treated with mouse IgG1 or 1 mM SmAb H1. Two-tailed unpaired t-test at time = 96 hours. B) Representative crystal violet staining of TC11 shCtrl and shSEMA7A KD#2 cells after treatment for 48 hours with mouse IgG1 or 1 mM SmAb H1. Bar graph: quantification of all replicates. Ordinary one-way ANOVA with Tukey’s multiple comparisons test. C) Average tumor volume in TC11 tumor-bearing mice over time, with groups receiving IgG1 or SmAb H1 (n=5 per group). Two-tailed unpaired t-test at day 4, 6, and 8 between (shCtrl) IgG and (shCtrl) SmAb H1 groups.. D) Tumor volume calculated using necropsy tumor photos in TC11 shCtrl and shSEMA7A-tumor bearing mice over time with groups receiving IgG1 or SmAb H1 (n=10 per group). Ordinary one-way ANOVA with Tukey’s multiple comparisons test. E) Weight of the tumor (and attached mammary gland) at study endpoint (n=10/group). Ordinary one-way ANOVA with Tukey’s multiple comparisons test. F) Immunohistochemistry staining of ex vivo TC11 tumors from IgG-treated and SmAb H1-treated groups. Scale bars = 50mm. Hematoxylin and eosin (H&E) staining (i-ii), active integrin B1 (9EG7) staining; brown (iii-iv), integrin B4 staining; brown (v-vi), and SEMA7A staining; brown (vii-viii). G) Log2(fold change) of pAKT (S473) expression in SmAb H1-treated versus IgG-treated TC11 tumors from RPPA-2 data. Two-tailed unpaired t-test. H) Quantification of 9EG7 (active ITGB1) IHC staining from all tumors (n=5 per group); Percent positive out of tumor area. Two-tailed unpaired t-test. I) Quantification of ITGB4 IHC staining from all tumors (n=5 per group); Percent positive out of tumor area. Two-tailed unpaired t-test. J) Quantification of SEMA7A IHC staining from all tumors (n=5 per group); percent positive out of tumor area. Two-tailed unpaired t-test. Error bars are mean +/- SD (A-B), or mean +/- SEM (C-E, G-I). *p<0.05, **p<0.01, ***p<0.001, ****p<0.0001.

### Combination treatment with fulvestrant and SmAbH1 reduces SEMA7A+ ER+ tumor growth

To test the hypothesis that the combination of SmAbH1 and fulvestrant may benefit patients, we tested cell viability in TC11 and MCF7 cell lines. TC11 shCtrl cell viability was inhibited by SmAbH1 alone, and even more so by combination of fulvestrant and SmAbH1, whereas TC11 shSEMA7A KD#2 viability was not affected (Figure 6A). Further, treatment of MCF7 EV and SEMA7A OE cells with fulvestrant, SmAbH1, or the combination showed that SEMA7A OE cells respond positively to SmAbH1 as a single agent and even more robustly to the combination of fulvestrant and SmAbH1, while control cells do not (SFigure 6A). Additionally, EV cells exhibit a moderate response to fulvestrant alone, while SEMA7A OE cells do not (SFigure 6A). These results were validated in SSM2 and ZR75-1 cells (SFigure 6B-C). In contrast, TC11 shSEMA7A cells displayed increased sensitivity to fulvestrant alone *in vitro* (Figure 6A), suggesting that loss of SEMA7A may re-sensitize TC11 tumor cells to fulvestrant. We also calculated synergy scores for fulvestrant in combination with SmAbH1 in TC11 cells (Figure 6B). SmAbH1 with fulvestrant resulted in a synergistic score (Bliss score>10), while SmAbH1 with tamoxifen was only slightly additive (Bliss score<10) (SFigure 6D), thus we moved forward with fulvestrant. To test our hypothesis *in vivo,* TC11 shCtrl and shSEMA7A KD#2 tumor cells were implanted into FVB/N mice and groups with Vehicle, IgG1, fulvestrant, SmAbH1, or the combination. Tumor growth of shCtrl tumors was reduced by SmAbH1 and the combination therapy, but not by the IgG or vehicle controls or fulvestrant alone (Figure 6C). Consistent with our hypothesis, growth of TC11 shSEMA7A tumors was not affected by SmAbH1 alone but was reduced by fulvestrant and the combination therapy (Figure 6D). Previous results were obtained with a hybridoma derived SmAbH1. Thus, we also tested a recombinant version of anti-SEMA7A antibody, rSmAbH1, for range of reactivity and specificity via IC50 and ELISA assay (SFigure 6E-F). Synergy results revealed that rSmAbH1-P and fulvestrant are synergistic *in vitro,* similar to what was observed with SmAbH1 and fulvestrant (SFigure 6G). We then repeated the TC11 *in vivo* experiment with rSmAbH1-P and fulvestrant; images of harvested tumors show visibly smaller tumors in the rSmAbH1-P-treated and combination-treated groups compared to controls or fulvestrant alone (Figure 6E). Mouse weights were monitored (SFigure 6H), and a 6-chemistry toxicity panel was run on mouse plasma and showed no differences between rSmAbH1-P-treated treatment and control-treated mice (SFigure 6I-N).

**Figure 6.**
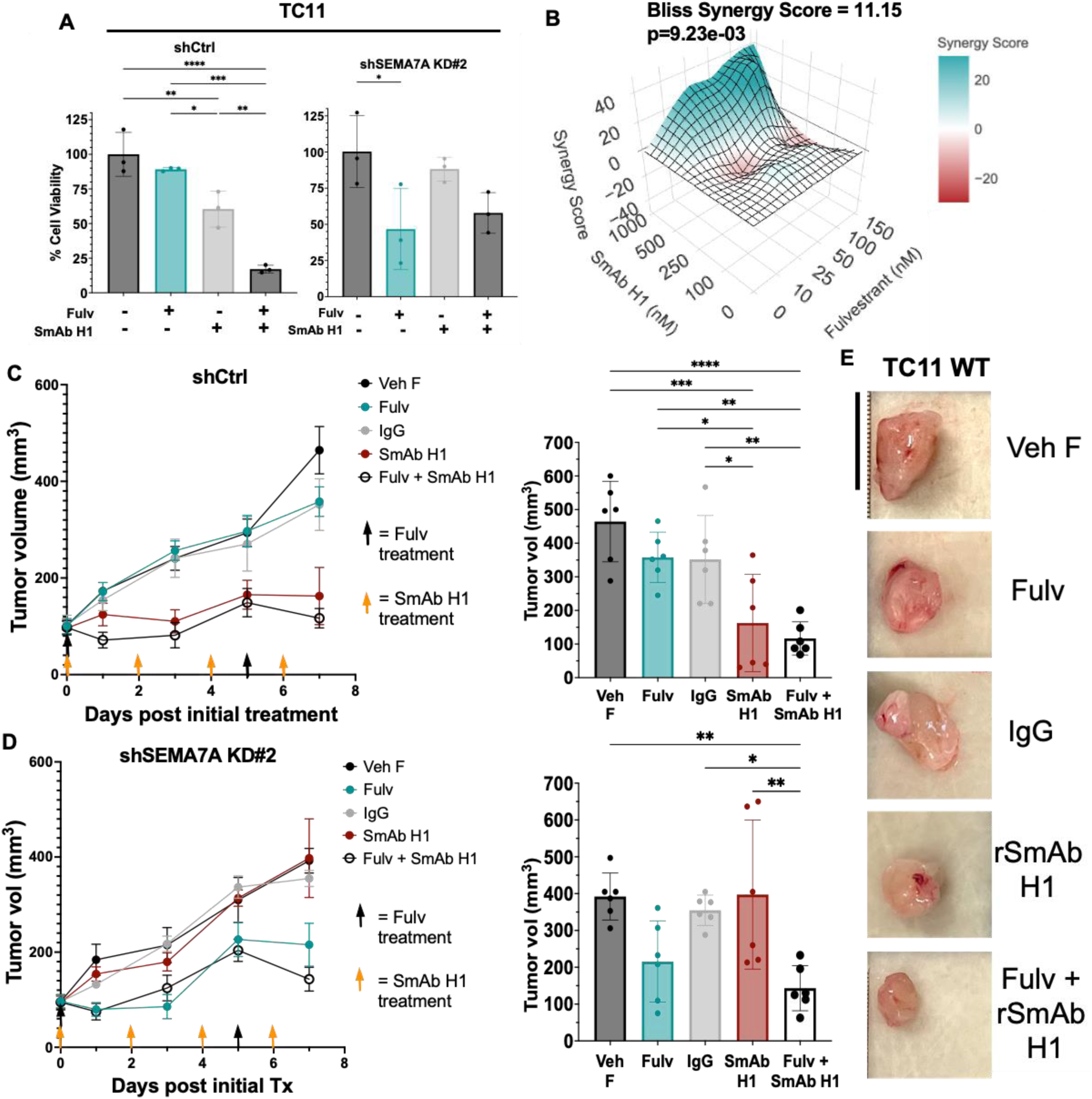
Combination treatment with fulvestrant and SmAb H1 reduces SEMA7A+ ER+ tumor growth. A) Cell viability quantified via crystal violet staining after 48-hour treatment of TC11 shCtrl and TC11 shSEMA7A KD#2 cells with DMSO, 8 nM fulvestrant, 1mM SmAb H1, or the combination, normalized to control. Ordinary one-way ANOVA with Tukey’s multiple comparisons test. B) Bliss synergy score for fulvestrant and SmAb H1 in TC11 cells. C) Average tumor volume in TC11 shCtrl tumor-bearing FVB/N mice over time with groups receiving Vehicle F, IgG1, fulvestrant, SmAb H1 or the combination (n=5 per group). Bar graph: tumor volume at study endpoint (Day 7). Ordinary one-way ANOVA with Tukey’s multiple comparisons test. D) Average tumor volume in TC11 shSEMA7A KD#2 tumor-bearing FVB/N mice over time with groups receiving Vehicle F, IgG1, fulvestrant, SmAb H1 or the combination (n=5 per group). Bar graph: tumor volume at study endpoint (Day 7). Ordinary one-way ANOVA with Tukey’s multiple comparisons test. E) Representative photographs of harvested TC11 tumors treated with Vehicle F, IgG1, fulvestrant, rSmAb H1 or the combination (n=5 per group). Scale bar = 1cm. Error bars are mean +/- SD (A), or mean +/- SEM (C-D). *p<0.05, **p<0.01, ***p<0.001, ****p<0.0001.

## DISCUSSION

Collectively, our results reveal mechanisms of action and therapeutic vulnerabilities of SEMA7A expressing ER+BC. Previous studies implicated integrin β1 as a receptor for SEMA7A on cancer cells and other cell types, our findings suggest that SEMA7A binds to both integrin β1 and β4 in ER+ breast cancer cells (Figure 2). We also provide evidence that the RGD motif within SEMA7A mediates binding to β-integrins. Furthermore, our findings with the RGDS blocking peptide suggest that SEMA7A enhances phosphorylation and activation of AKT specifically through integrins, which were previously shown to undergo conformational changes when bound by an RGD motif leading to intracellular activation of signaling pathways such as PI3K/AKT, ILK, Fak, and MAPK/ERK (49). Our results are consistent with previously results showing that integrin β1 activation of PI3K and MAPK signaling can facilitate endocrine therapy resistance in breast cancer (50); thus blocking this SEMA7A mediated pathway is a potential way to overcome endocrine therapy resistance in patients with ER+BC. Limitations of the current study include that the synthetic RGDS peptide may have also caused non-specific integrin inhibition beyond that attributed to SEMA7A. Further, while the RGD motif in SEMA7A is known, the protein folding structure of SEMA7A when bound versus unbound to integrins remains unknown. Future studies could further investigate this protein interaction through mutations and further structural analyses.

Our findings provide premise for treating patients with SEMA7A+ breast cancer with more targeted therapies, including those that target of PI3K and/or directly target SEMA7A. Given our previous studies showing poor relapse-free survival rates in both ER+ and ER-breast cancer patients (2), clinical trials informed by our data stand to impact thousands of patients annually. We observed a correlation between SEMA7A overexpression and increased pAKT (S473) and downstream targets of AKT, suggesting activation of downstream survival signaling by AKT. Since ILK was moderately increased in SEMA7A OE cells and ILK is known to phosphorylate AKT at S473, we suspect involvement of ILK downstream of SEMA7A/integrin signaling. Because there are no clinically available integrin or ILK inhibitors, we suggest that clinically approved PI3K inhibitors be tested to inhibit the SEMA7A-AKT pathway. Clinical use of alpelisib is currently approved for patients with an E545K mutation in *PIK3CA* that causes constitutive activation of the PI3K p110α subunit (24,51).

However, we hypothesize that even in the absence of the *PIK3CA* E545K mutation, high SEMA7A expression can cause hyperactivation of PI3K including p110α, thus SEMA7A-expressing tumors may be sensitive to alpelisib (52). GCT-007, a pan-PI3K inhibitor that has shown preclinical efficacy in glioblastoma, is not approved yet for clinical use, but our results suggest it may be efficacious in breast cancer patients as well. Limitations of the clinical applicability of our study include the known toxicities observed in some patients including hyperglycemia, rash, and diarrhea (53), and while we saw no overt toxicities in our mouse studies, the potential toxicities of GCT-007 in humans is unknown. Additionally, while we did not explore direct targeting of AKT future studies could determine that this is another viable option for patients with ER+SEMA7A+BC. Pan-AKT inhibitors such as capivasertib and ipatasertib, mTOR inhibitors such as everolimus, and other PI3K inhibitors such as a p110δ inhibitors (idelalisib) could be explored in our SEMA7A+ER+BC models to optimize available treatment combinations. Moreover, we have published that the Bcl-2 inhibitor venetoclax can slow tumor growth in pre-clinical models, and our RPPA-1 results confirm increased Bcl-2 expression with SEMA7A expression; thus, combination of SmAbH1 and venetoclax should also be explored.

Our results suggest that direct inhibition of SEMA7A via our monoclonal antibody, SmAbH1, is a promising therapeutic strategy that should be further investigated with additional pre-clinical studies to enable progression to first in human clinical trials. While the combination of fulvestrant and SmAbH1 did reach statistical significance compared to SmAbH1 alone, it is important to note that SmAbH1 did not appear to interfere with fulvestrant, which is utilized clinically for ER+ breast cancers and at study end the tumors were visibly smaller in the combination-treated group. These results suggest that patients will obtain benefit from the addition of SmAbH1 to their current standard of care (continued endocrine therapy) in the recurrent/metastatic setting and that after additional testing SmAbH1 could become a novel front line therapy for ER+BC patients that express SEMA7A. Aside from reduced tumor growth, we also saw that SmAbH1 decreased pAKT (S473) suggesting that SmAbH1 is functionally inhibiting the pathway activated by SEMA7A/integrin interaction. We also performed additional in vivo SmAb/fulvestrant studies with a newly generated recombinant antibody, rSmAbH1-P, which is necessary for humanization and future IND-enabling studies. A limitation of our study is the use of mouse cell lines for the *in vivo* models instead of human cell lines. We deliberately chose the TC11 and SSM2 models to test SmAbH1 because they are immunocompetent and we have recently published that SmAbH1 activates anti-tumor T cell responses by re-activating anti-tumor T cell responses (29).

We conclude that inhibition of SEMA7A mediated signaling in breast cancer, both direct and indirect, are promising clinical treatment avenues that should be further investigated particularly in patients with recurrent disease despite endocrine therapies. Furthermore, our previous results suggest that SEMA7A positive tumors will not respond to CDK4/6 inhibition, which is FDA approved for ER+ breast cancer patients. Additionally, our results suggest that patients with ER+ BC may benefit from a diagnostic test that quantifies SEMA7A expression as we have published a threshold of SEMA7A expression by immunostaining fixed tissues that can predict for relapse in a small patient dataset(17). If such a diagnostic existed, patients with SEMA7A+ tumors could receive a SEMA7A-targeted therapy like patients with HER2+ disease that are treated with trastuzumab. Future directions could also include the development and optimization of a minimally invasive SEMA7A-detecting diagnostic (blood) test, which we have initiated. However, further investigation is needed to determine whether SEMA7A levels in tumors correlate with levels in patient serum samples. Cumulatively, our current and previous studies suggest a unique and druggable aspect in the biology of SEMA7A-expressing breast cancers that could transform treatment options for ER+BC.

## ACKNOWLEDGEMENTS

Technical support

`o` Lyons Lab: Lyndsey Crump, Lauren Cozzens, Kelsey Kines, Alan Elder, Alex Becks (University of Colorado Anschutz)
`o` Sarah Tarullo (Plymouth State University)
`o` Porter lab: Weston Porter, Garhett Wyatt (Texas A&M)
- Provided Reagents:

`o` Global Cancer Technology
`o` Novartis
- Clinical mentor:

- Virginia Borges (CU Anschutz, UC Health)
- Thesis committee members: Carol Sartorius, Benjamin Bitler, Katherine Fantauzzo, John Tentler, Jennifer Richer, Daniel Sherbenou
- Experimental advice:

- Michael Oliphant
- NIH/NCATS Colorado CTSA Grant Number TL1 TR002533 to RS
- NIC R01CA282900-01 and R01HD108335-01A1 to TL
- SPARK/REACH Program, NIH: U01HL152405 and OEIDIT/AIA: MA 2021-2028 to TL
- ABN Ohana Gates Grubstake Award to TL
- DOM Outstanding Early Career Scholars Program to TL
- UCCC and DOM ASPIRE to TL
- Men for the Cure PO1 pilot grant to TL
- University of Colorado Cancer Center Support Grant #P30CA046934 to RS

`o` Anh Le, Steve Anderson and the University of Colorado Anschutz CTSR (Cell Technologies Shared Resource)
`o` University of Colorado Anschutz CPSR (Comparative Pathology Shared Resource)
- Cancer Prevention & Research Institute of Texas Core Facility Support Awards (PR210227, PI: Dean Edwards)

- Baylor College of Medicine Antibody-Based Proteomics Core
- Dean Edwards
- Shixia Huang

