## Supplemental Data for "Targeting Semaphorin 7a signaling in preclinical models of estrogen receptor-positive breast cancer"

**Supplemental Figure 1.**

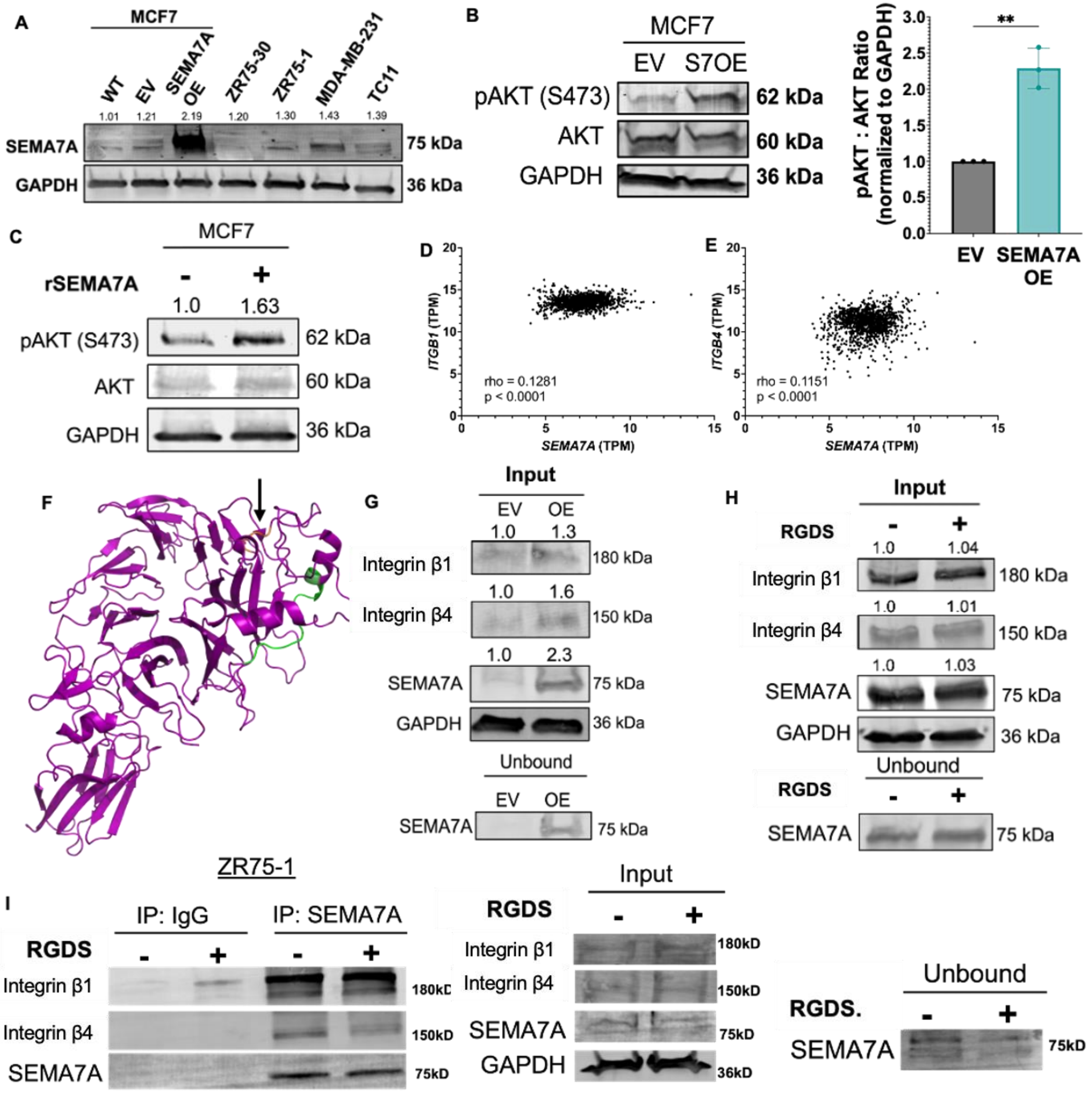

**Supplemental Figure 1 Legend:**

A) Representative immunoblot for SEMA7A and GAPDH in breast cancer cell lines. (WT = wild type, EV = empty vector lentiviral expression, OE = lentiviral SEMA7A overexpression.

B) Representative immunoblot for pAKT (S473), AKT and GAPDH in MCF7 EV and SEMA7A OE cell lines. Bar graph: ratio of pAKT to total AKT. Two-tailed unpaired t-test.

C) Representative immunoblot for pAKT (S473), AKT and GAPDH in MCF7 wild type cells after treatment for 1 hour with 20 μg/mL rSEMA7A or control (PBS).

D) Correlation of *SEMA7A* and *ITGB1* mRNA transcripts per million (TPM), from TCGA Breast Cancer Dataset (n=1247), log2(norm+1).

E) Correlation of *SEMA7A* and *ITGB4* mRNA transcripts per million (TPM), from TCGA Breast Cancer Dataset (n=1247), log2(norm+1).

F) Ribbon structure of SEMA7A protein with RGD motif in orange (aa 267-269) and the site at which it binds to the monoclonal antibody (SmAb H1) in green (aa 381-392).

G) CoIP Input: Immunoblot for integrin β1, integrin β4, SEMA7A and GAPDH in MCF7 EV or SEMA7A OE lysates, and for SEMA7A in the unbound sample.

H) CoIP Input: Immunoblot for integrin β1, integrin β4, SEMA7A, and GAPDH in MCF7 SEMA7A OE lysates after treatment with PBS or 50 μM RGDS peptide for 1 hour, and for SEMA7A in the unbound sample.

I) Left: Immunoblot integrin β1, integrin β4, and SEMA7A in elutions following CoIP of lysates for IgG or SEMA7A, after treatment with PBS (-) or 50 μM RGDS (+). Middle: Immunoblot for ITGB4, ITGB1, SEMA7A, and GAPDH in ZR751 cell lysates after treatment with PBS (-) or RGDS peptide (+) for 1 hour (input). Right: Immunoblot for SEMA7A in unbound sample following IP.

Error bars are mean +/- SD. *p<0.05, **p<0.01, ***p<0.001, ****p<0.0001.

**Supplemental Figure 2.**

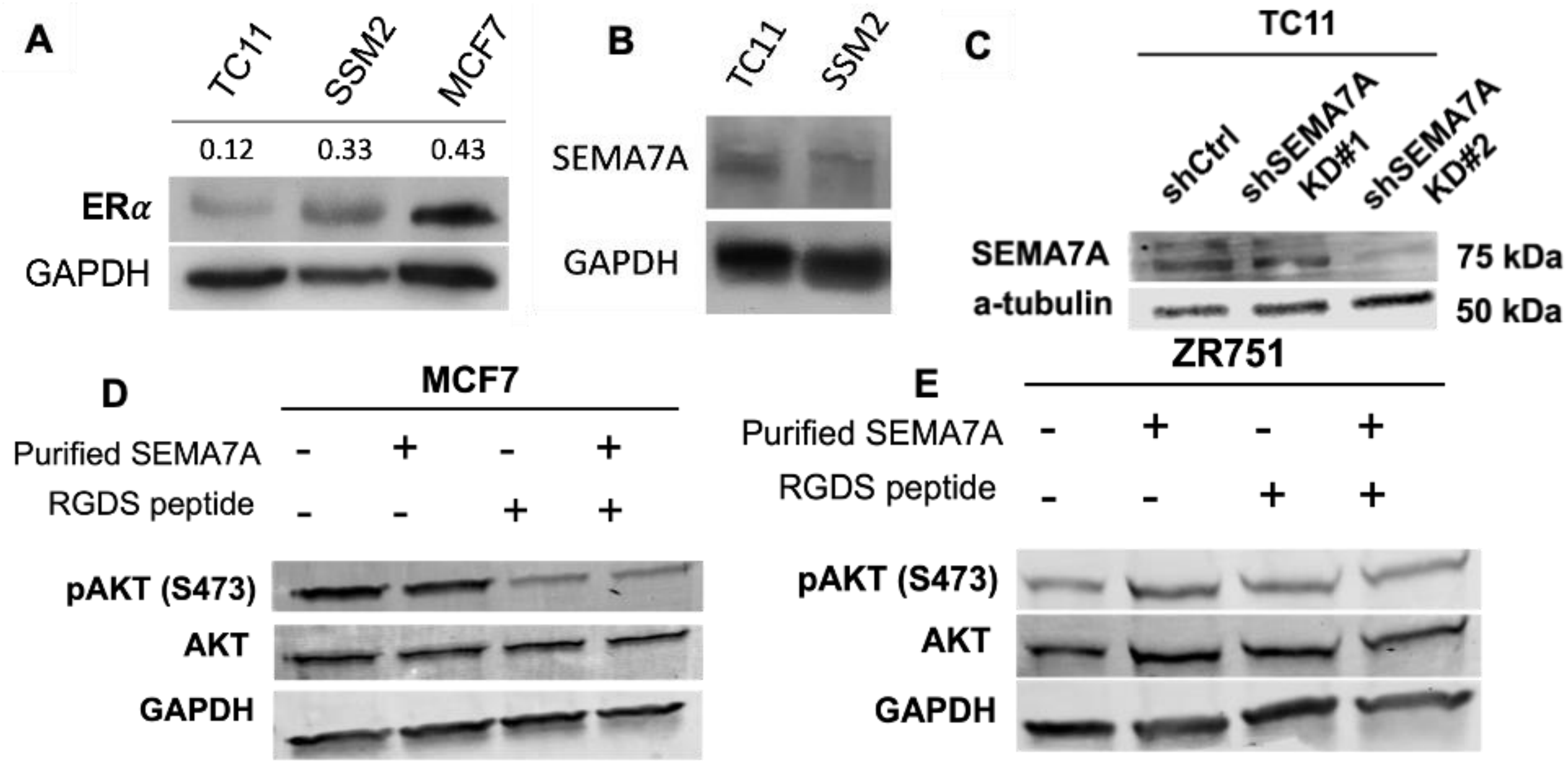

**Supplemental Figure 2 Legend:**

A) Representative immunoblot for ERα and GAPDH in TC11, SSM2 and MCF7 cell line lysates.

B) Representative immunoblot for SEMA7A and GAPDH in TC11 and SSM2 cell line lysates.

C) Representative immunoblot for SEMA7A and GAPDH in TC11 shCtrl, TC11 shSEMA7A KD#1 and TC11 shSEMA7A KD#2 cell lysates.

D) Representative immunoblot for pAKT (S473), AKT and GAPDH in MCF7 WT cell lysates after treatment for 1 hour with rSEMA7A, RGDS peptide, or both.

E) Representative immunoblot for pAKT (S473), AKT and GAPDH in ZR75-1 cell lysates after treatment for 1 hour with rSEMA7A protein, RGDS peptide, or both.

**Supplemental Figure 3.**

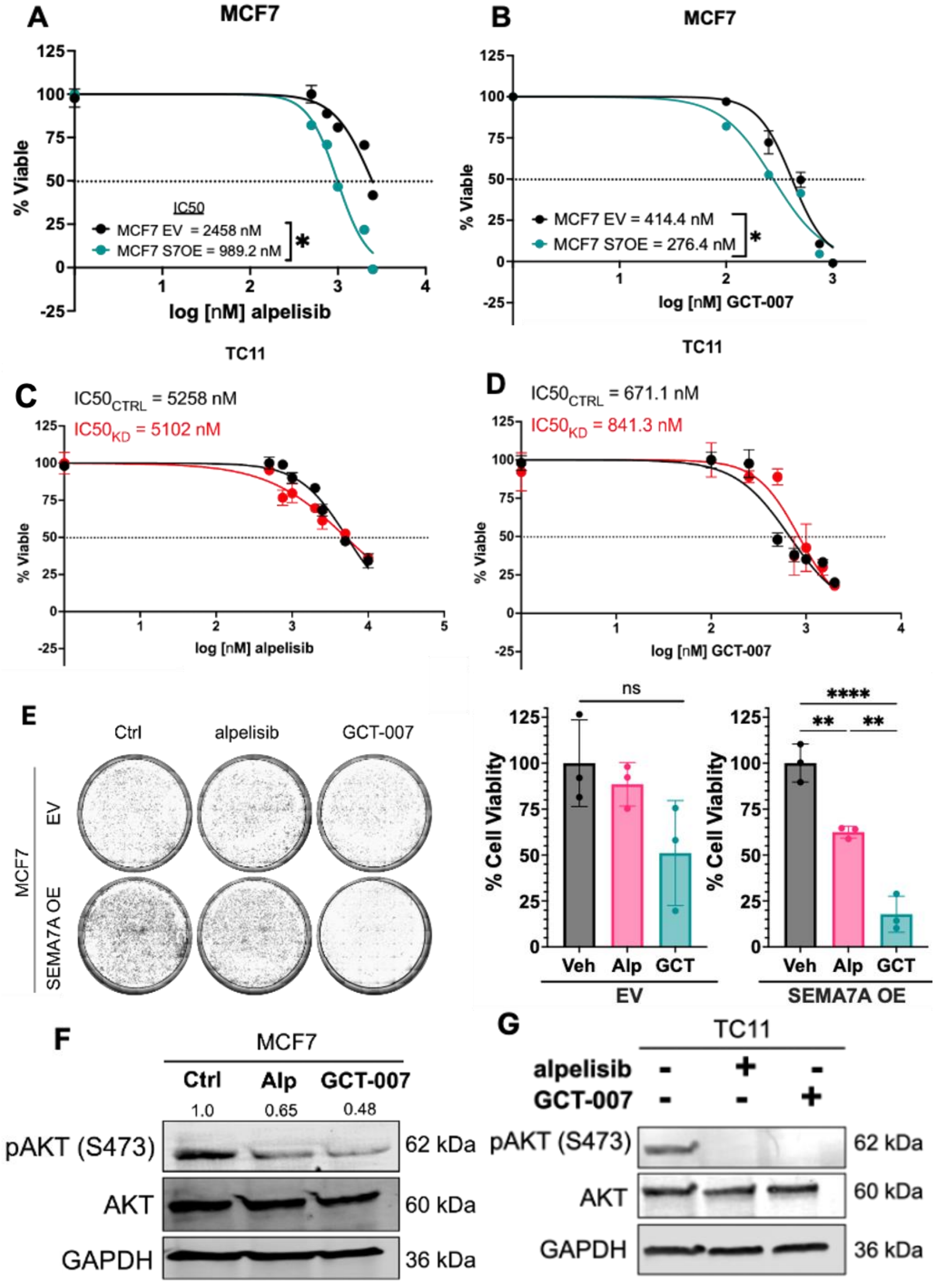

**Supplemental Figure 3 Legend:**

A) IC50 dose curve generated by treating MCF7 EV and SEMA7A OE cells with increasing doses of alpelisib (0, 0.5, 0.75, 1.0, 2.0, and 2.5 μM) for 48 hours.

B) IC50 dose curve generated by treating MCF7 EV and SEMA7A OE cells with increasing doses of GCT-007 (0, 0.1, 0.25, 0.5, 0.75, and 1.0 μM) for 48 hours.

C) IC50 dose curve generated by treating TC11 shCtrl and shSEMA7A#2 cells with increasing doses of alpelisib (0, 0.5, 0.75, 1.0, 2.0, and 2.5 μM) for 48 hours.

D) IC50 dose curve generated by treating TC11 shCtrl and shSEMA7A#2 cells with increasing doses of GCT-007 (0, 0.1, 0.25, 0.5, 0.75, and 1.0 μM) for 48 hours.

E) Representative results from cell viability assay: MCF7 EV and SEMA7A OE cells were treated with 1 μM DMSO, 1 μM alpelisib, or 0.3 μM GCT-007 for 48 hours, followed by crystal violet staining. Bar graph: quantification of cell viability using ImageJ analysis. Standard one-way ANOVA with Tukey's multiple comparisons test.

F) Representative immunoblot for pAKT (S473), AKT and GAPDH, of MCF7 cell lysates after 48 hour treatment with 1 μM DMSO, 1 μM alpelisib, or 0.3 μM GCT-007.

G) Representative immunoblot for pAKT (S473), AKT and GAPDH, of TC11 cell lysates after 48-hour treatment with 1 μM DMSO, 1 μM alpelisib, or 0.3 μM GCT-007.

Error bars are mean +/- SD. *p<0.05, **p<0.01, ***p<0.001, ****p<0.0001.

**Supplemental Figure 4.**

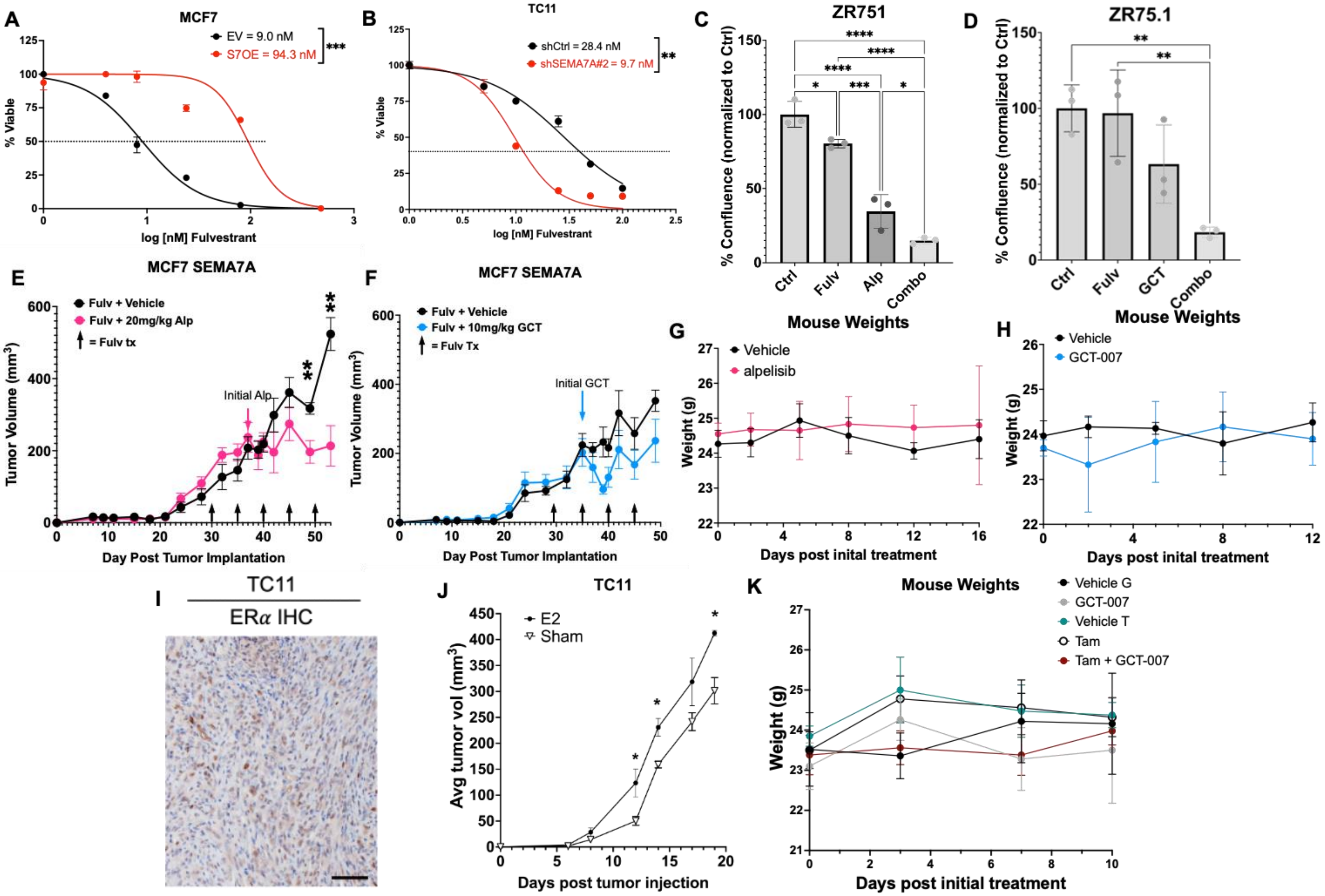

**Supplemental Figure 4 Legend:**

A) IC50 dose curve generated by treating MCF7 EV and SEMA7A OE cells with increasing doses of fulvestrant (0, 4, 8, 24, 80, 240 and 480 nM) for 48 hours.

B) IC50 dose curve generated by treating TC11 shCtrl and shSEMA7A#2 cells with increasing doses of fulvestrant (0, 4, 8, 24, 80, 240 and 480 nM) for 48 hours.

C) Cell viability quantified as cell confluence normalized to control, via crystal violet staining after 48-hour treatment of ZR75-1 cells with DMSO, 8nM fulvestrant, 1µM alpelisib, or the combination.

D) Cell viability quantified as cell confluence normalized to control, via crystal violet staining after 48-hour treatment of ZR75-1 cells with DMSO, 8nM fulvestrant, 0.3µM GCT-007, or the combination.

E) Average MCF7 SEMA7A OE tumor volume in NCG mice over time, treated with either fulvestrant alone or the combination of fulvestrant plus alpelisib (n=5 per group). Unpaired two-tailed T test at each measured timepoint.

F) Average MCF7 SEMA7A OE tumor volume in NCG mice over time, treated with either fulvestrant alone or the combination of fulvestrant plus GCT-007 (n=5 per group). Unpaired two-tailed T test at each measured timepoint.

G) MCF7 SEMA7A OE-tumor bearing NCG mouse weights over time after treatment with vehicle or alpelisib.

H) MCF7 SEMA7A OE-tumor bearing NCG mouse weights over time after treatment with vehicle or GCT-007.

I) TC11 tumor IHC stained for estrogen receptor alpha; brown. Scale bars = 30µm.

J) TC11 average tumor volume in FVB/N mice over time, treated with sham or estrogen (E2) pellets.

K) TC11-tumor bearing FVB/N mouse weights over time after treatment with Vehicle G, GCT-007, Vehicle T, Tamoxifen, or the combination of Tamoxifen and GCT-007.

Error bars are mean +/- SD (C-D), or mean +/- SEM (E-J). *$p<0.05$, **$p<0.01$, ***$p<0.001$, ****$p<0.0001$.

**Supplemental Figure 5.**

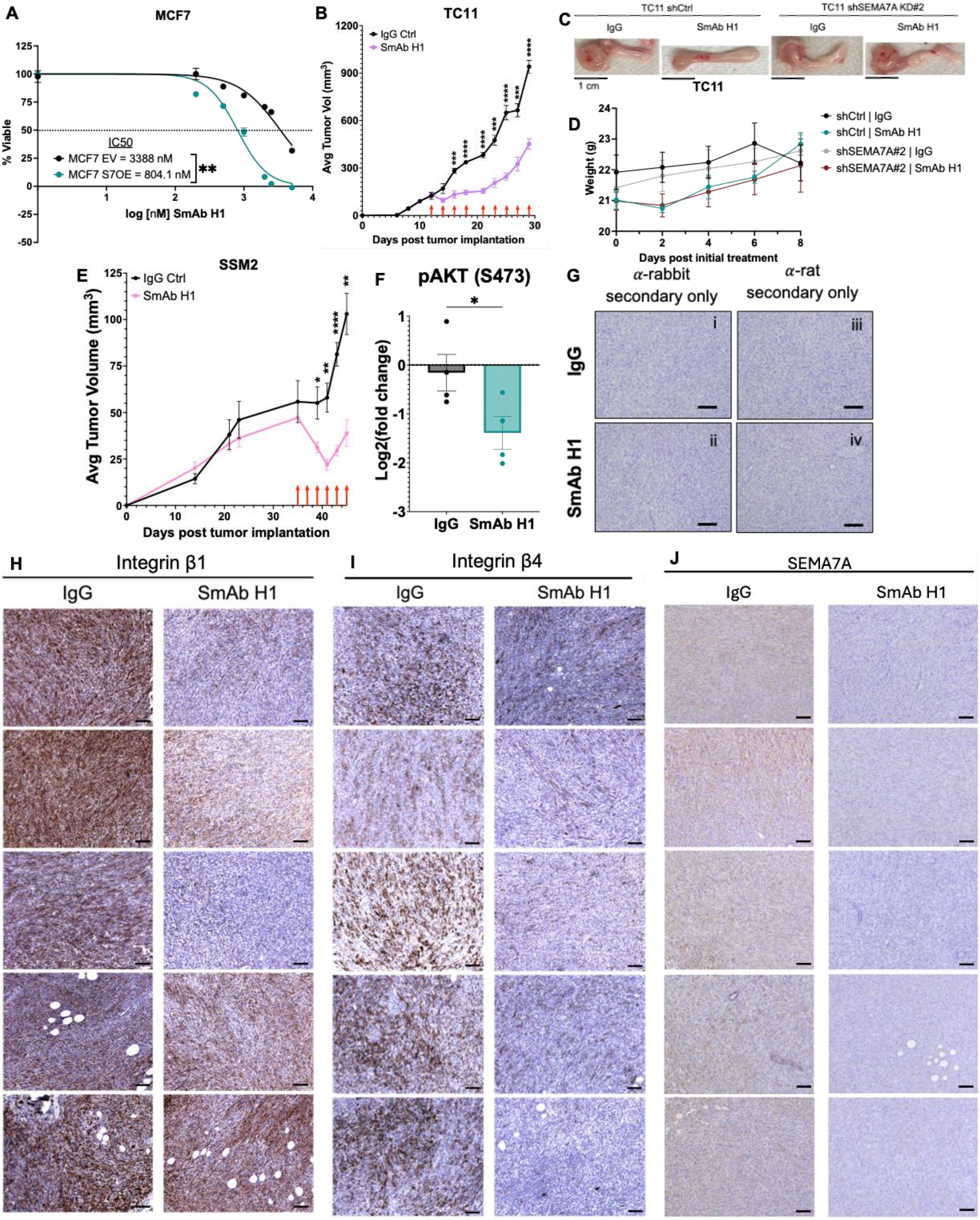

**Supplemental Figure 5 Legend:**

A) IC50 dose curve generated by treating MCF7 EV and SEMA7A OE cells with increasing doses of SmAb H1 (0, 200, 500, 1000, 2000, 2500, and 5000 nM) for 48 hours.

B) Average tumor volume of TC11 tumor-bearing FVB/N mice over time, during treatment with either IgG or SmAb H1 (n=5/group). Unpaired two-tailed T test at each measured timepoint.

C) Representative photographs of harvested TC11 shCtrl and shSEMA7A#2 tumors treated with IgG1 or SmAb H1 (n=5 per group). Scale bar = 1cm.

D) TC11 shCtrl and shSEMA7A#2-tumor bearing FVB/N mouse weights over time after treatment with IgG1 or SmAb H1.

E) Average tumor volume of SSM2 tumor-bearing 129SV-E mice over time, during treatment with either IgG or SmAb H1 (n=5/group). Unpaired two-tailed T test at each measured timepoint.

F) Log2(fold change) of pAKT (S473) expression in SmAb H1-treated versus IgG-treated SSM2 tumors from RPPA-2 data. Two-tailed unpaired t-test.

G) IHC negative controls; TC11 IgG or SmAb H1-treated tumors stained with secondary antibody only (anti-rabbit and anti-rat).

H) TC11 IgG and SmAb H1-treated tumors IHC stained for active integrin β1 (9EG7); brown. Scale bars = 50μm.

I) TC11 IgG and SmAb H1-treated tumors IHC stained for integrin β4; brown. Scale bars = 50μm.

J) TC11 IgG and SmAb H1-treated tumors IHC stained for SEMA7A; brown. Scale bars = 50μm.

Error bars are mean +/- SEM. *p<0.05, **p<0.01, ***p<0.001, ****p<0.0001.

**Supplemental Figure 6.**

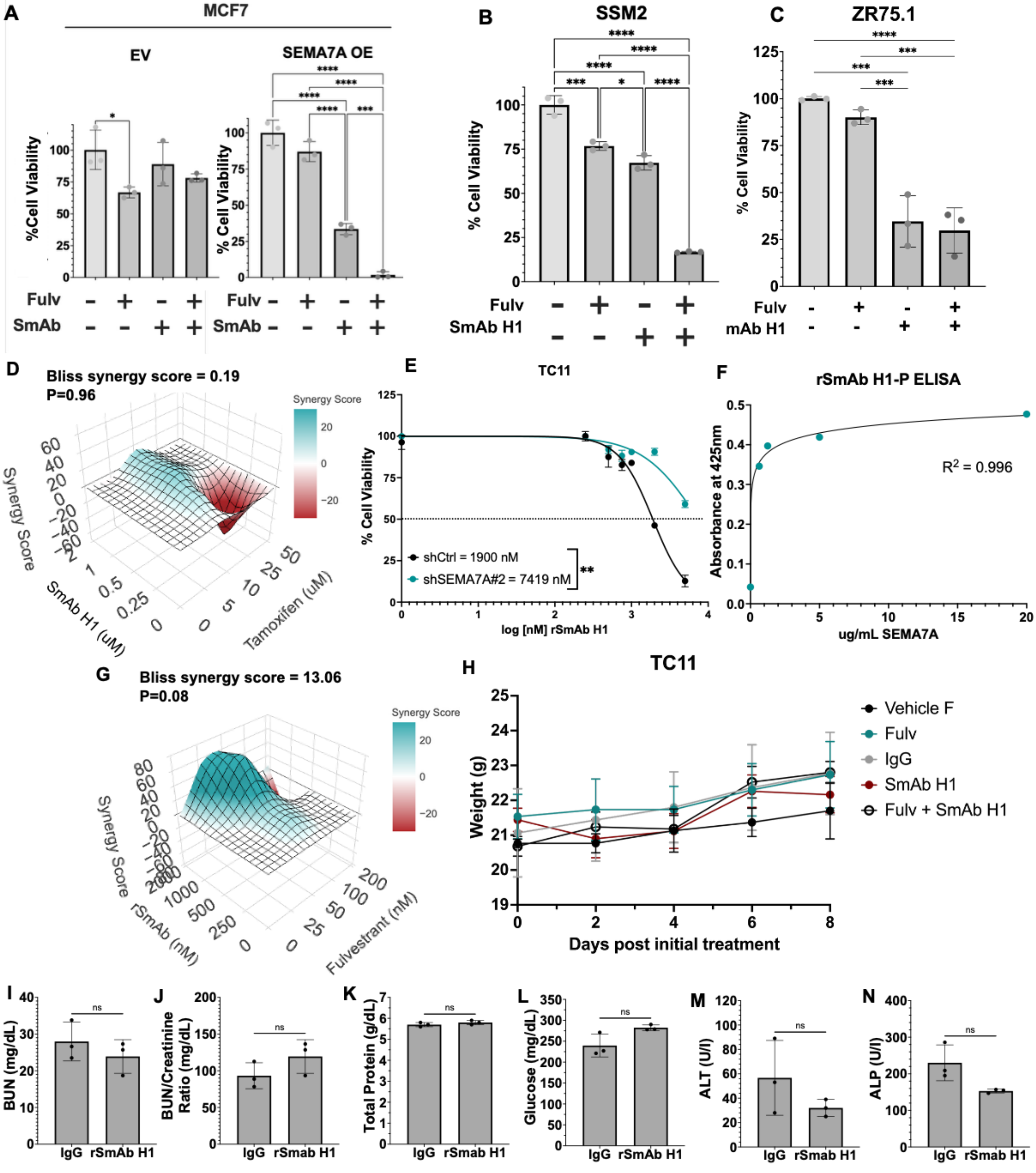

**Supplemental Figure 6 Legend:**

A) Cell viability quantified via crystal violet staining after 48-hour treatment of MCF7 EV and SEMA7A OE cells with DMSO, fulvestrant, SmAb H1, or the combination. Standard one-way ANOVA with Tukey's multiple comparisons test.

B) Cell viability quantified via crystal violet staining after 48-hour treatment of SSM2 cells with DMSO, fulvestrant, SmAb H1, or the combination. Standard one-way ANOVA with Tukey's multiple comparisons test.

C) Cell viability quantified via crystal violet staining after 48-hour treatment of ZR75.1 cells with DMSO, fulvestrant, SmAb H1, or the combination. Standard one-way ANOVA with Tukey's multiple comparisons test.

D) Bliss synergy scores for tamoxifen and SmAb H1 in TC11 cells.

E) IC50 dose curve generated by treating TC11 shCtrl and shSEMA7A#2 cells with increasing doses of rSmAb H1-P for 48 hours.

F) Indirect ELISA performed using increasing concentrations of SEMA7A protein-coated wells and rSmAb H1-P as the primary antibody.

G) Bliss synergy scores for fulvestrant and rSmAb H1-P in TC11 cells.

H) TC11 -tumor bearing FVB/N mouse weights over time after treatment with Vehicle F, fulvestrant, IgG1, SmAb H1, or the combination of fulvestrant and SmAb H1.

I) Blood urea nitrogen (BUN) levels (mg/dL) in mouse serum from IgG and rSmAb H1-P treated TC11 mice.

J) Blood urea nitrogen (BUN) to Creatinine Ratio in mouse serum from IgG and rSmAb H1-P treated TC11 mice.

K) Total protein levels (g/dL) in mouse serum from IgG and rSmAb H1-P treated TC11 mice.

L) Glucose levels (mg/dL) in mouse serum from IgG and rSmAb H1-P treated TC11 mice.

M) Alanine aminotransferase (ALT) levels (U/I) in mouse serum from IgG and rSmAb H1-P treated TC11 mice.

N) Alkaline phosphatase (ALP) levels (U/I) in mouse serum from IgG and rSmAb H1-P treated TC11 mice.

Error bars are mean +/- SD (A-C), or mean +/- SEM (I-N). *p<0.05, **p<0.01, ***p<0.001, ****p<0.0001.
